# Null effects of levodopa on reward- and error-based motor adaptation, savings, and anterograde interference

**DOI:** 10.1101/2020.11.19.390302

**Authors:** Dimitrios J. Palidis, Heather R. McGregor, Andrew Vo, Penny A. MacDonald, Paul L. Gribble

**Affiliations:** The Brain and Mind Institute, Western University, London, ON, N6A 57B, Canada; Dept. Psychology, Western University, London, ON, N6A 5C2, Canada; Graduate Program in Neuroscience, Schulich School of Medicine & Dentistry, Western University, London, ON, N6A 3K7, Canada; Dept. of Applied Physiology and Kinesiology, University of Florida, Gainesville, FL, USA; Dept. of Neurology and Neurosurgery, Montreal Neurological Institute, McGill University, Montreal, QC, H3A 2B4, Canada; Dept. Physiology & Pharmacology, Schulich School of Medicine & Dentistry, Western University, London, ON, N6A 5C1 Canada; Dept. of Clinical Neurological Sciences, University of Western Ontario, London, ON, N6A 5A5, Canada; Haskins Laboratories, New Haven, CT, 06511, USA

**Keywords:** motor adaptation, reward, reinforcement, FRN, reward positivity, dopamine

## Abstract

Dopamine signaling is thought to mediate reward-based learning. We tested for a role of dopamine in motor adaptation by administering the dopamine precursor levodopa to healthy participants in two experiments involving reaching movements. Levodopa has been shown to impair reward-based learning in cognitive tasks. Thus, we hypothesized that levodopa would selectively impair aspects of motor adaptation that depend on reinforcement of rewarding actions.

In the first experiment, participants performed two separate tasks in which adaptation was driven either by visual error-based feedback of the hand position or binary reward feedback. We used EEG to measure event-related potentials evoked by task feedback. We hypothesized that levodopa would specifically diminish adaptation and the neural responses to feedback in the reward learning task. However, levodopa did not affect motor adaptation in either task nor did it diminish event-related potentials elicited by reward outcomes.

In the second experiment, participants learned to compensate for mechanical force field perturbations applied to the hand during reaching. Previous exposure to a particular force field can result in savings during subsequent adaptation to the same force field or interference during adaptation to an opposite force field. We hypothesized that levodopa would diminish savings and anterograde interference, as previous work suggests that these phenomena result from a reinforcement learning process. However, we found no reliable effects of levodopa.

These results suggest that reward-based motor adaptation, savings, and interference may not depend on the same dopaminergic mechanisms that have been shown to be disrupted by levodopa during various cognitive tasks.

**New and Noteworthy:** Motor adaptation relies on multiple processes including reinforcement of successful actions. Cognitive reinforcement learning is impaired by levodopa-induced disruption of dopamine function. We administered levodopa to healthy adults who participated in multiple motor adaptation tasks. We found no effects of levodopa on any component of motor adaptation. This suggests that motor adaptation may not depend on the same dopaminergic mechanisms as cognitive forms or reinforcement learning that have been shown to be impaired by levodopa.

## Introduction

Human motor control is adaptive to changes of the environment and the body through multiple mechanisms including reinforcement of successful actions and recalibration of internal mappings between motor commands and sensory outcomes (Huang et al., 2011; Izawa & Shadmehr, 2011; J. A. Taylor et al., 2014; Wolpert et al., 1995). Two prominent experimental models of motor adaptation are force field adaptation and visuomotor rotation (VMR) tasks. In studies of force field adaptation, a robot applies velocity-dependent forces to the hand during reaches to targets. In visuomotor rotation tasks, a cursor on a digital display represents the position of the hand, and the mapping between the actual reach angle and the position of the cursor is rotated. In both tasks participants quickly adapt their movements to compensate for the experimentally induced perturbations. Learning involves the cerebellum, and parietal, sensory, and motor cortical areas (Diedrichsen et al., 2005; Ito, 2000; Krakauer et al., 2004; Smith & Shadmehr, 2005; Tanaka et al., 2009; Jordan A. Taylor et al., 2010; Wong et al., 2019). It is thought that these neural circuits predict the sensory consequences of motor commands, and that adaptation occurs in response to sensory prediction error when sensory afference violates these predictions (Adams et al., 2013; Bhanpuri et al., 2013; Izawa & Shadmehr, 2011; R. Chris Miall et al., 2007; Shadmehr et al., 2010; Synofzik et al., 2008; Therrien & Bastian, 2015; Tseng et al., 2007; Wolpert et al., 1995).

While sensory error-based learning mechanisms are dominant in typical motor adaptation paradigms, influences of reinforcement learning processes are increasingly recognized (Bernardi et al., 2015; Cashaback et al., 2019; Izawa & Shadmehr, 2011; Kim et al., 2019; Kooij et al., 2018; McDougle et al., 2016; Mehler et al., 2017; Nikooyan & Ahmed, 2014; Palidis et al., 2019; Sidarta et al., 2016, 2018; van der Kooij & Smeets, 2019). Reward and task success can modulate sensory error-based learning (Galea et al., 2015; Kim et al., 2019; Kooij et al., 2018; Kuling et al., 2019; Leow et al., 2018, 2020; Shmuelof et al., 2012). Reinforcement learning and sensory error-based learning can also contribute to adaptation as separable processes. Adaptation to sensory error has been shown to occur automatically even when it interferes with task success (Mazzoni & Krakauer, 2006). Reward-based adaptation can be isolated experimentally by providing only binary reinforcement feedback indicating success or failure (Izawa & Shadmehr, 2011; Shmuelof et al., 2012). When sensory error-based learning cannot occur due to impoverished sensory feedback or cerebellar damage, reward-based learning can produce comparable behavioral adaptation (Cashaback et al., 2017; Izawa & Shadmehr, 2011; Therrien et al., 2016).

It is thought that reward prediction error drives biological reinforcement learning when an action results in an outcome that is better or worse than expected (Daw & Tobler, 2014; Sambrook & Goslin, 2015; Walsh & Anderson, 2012). Phasic changes in the firing rate of midbrain dopamine neurons match reward prediction error signals predicted by computational models of reinforcement learning (Bayer & Glimcher, 2005; García-García et al., 2017; Jocham & Ullsperger, 2009; Schultz et al., 1997; Watabe-Uchida et al., 2017). These dopaminergic signals are thought to mediate synaptic plasticity in the striatum and frontal cortex underlying reward-based learning (Otani et al., 2003; Reynolds & Wickens, 2002; Wang et al., 2018).

Levodopa is a dopamine precursor commonly used to treat motor symptoms in patients with Parkinson’s disease. Levodopa has been shown to impair reward-based learning in both patients and healthy participants (Cools et al., 2001, 2007; Feigin et al., 2003; Frank et al., 2004; Graef et al., 2010; Hiebert et al., 2014; Jahanshahi et al., 2010; Kwak et al., 2010; MacDonald et al., 2011; Swainson et al., 2000; Torta et al., 2009; Vo et al., 2016, 2018). According to the “dopamine overdose” hypothesis, dopamine levels affect performance in tasks that depend on the striatum according to an inverted-u function (Cools et al., 2007). In early-stage Parkinson’s disease, the dorsal striatum is significantly depleted of dopamine whereas the ventral striatum is comparatively spared. Dopaminergic therapy is predicted to ameliorate deficits caused by dopamine-depletion in the dorsal striatum but to worsen functions ascribed to the ventral striatum. In line with this view, reward-based learning is thought to rely on dopamine signaling in ventral striatum and is impaired by levodopa.

Although dopamine is widely implicated in reward-based learning, it is not clear whether this role extends to reward-based motor adaptation. We administered levodopa to healthy young participants to test for effects on motor adaptation. In our first experiment, participants received levodopa and placebo in separate sessions using a repeated measures design. Both sessions included a reward-based learning task and a sensory error-based VMR task. In the reward-based learning task, adaptation was induced through binary reinforcement feedback at the end of each movement. We measured changes in the mean reach angle due to reinforcement as well as modulations in trial-by-trial variability of reach angle as a response to reward outcomes. Previous research has shown that motor variability increases following unrewarded outcomes compared to rewarded outcomes (Dhawale et al., 2019; Holland et al., 2018; Mastrigt et al., 2020; Pekny et al., 2015; van der Kooij & Smeets, 2019). This could indicate reinforcement of rewarded actions as well as exploration in response to unrewarded outcomes (Cashaback et al., 2019; Dhawale et al., 2019). This variance modulation is impaired in individuals with Parkinson’s disease who are medicated, but it remains unclear whether this deficit is caused by the disease process itself or side-effects of dopaminergic medication (Pekny et al., 2015). We predicted that levodopa would impair reward-based motor adaptation and modulation of trial-by-trial variability in accordance with the “dopamine overdose hypothesis”.

In the sensory error-based learning task, participants adapted to visuomotor rotation perturbations designed to produce sensory prediction error while minimizing reward prediction error. Investigations as to whether VMR learning depends on dopamine have shown inconsistent results (Bédard & Sanes, 2011; Marinelli et al., 2009; Mongeon et al., 2013; Noohi et al., 2014). Sensory error-based learning may be mediated by non-dopaminergic mechanisms depending primarily on the cerebellum, whereas dopamine affects VMR learning through additional or modulatory contributions of a reinforcement learning process (Singh et al., 2019). We hypothesized that sensory error-based learning would be unaffected by levodopa. As such, we designed our sensory error-based learning task to preclude effects of reinforcement.

In experiment 1, we recorded EEG to measure neural event-related potentials (ERPs). Previously, we found that a medial frontal ERP component called the feedback related negativity, or alternatively the reward positivity (FRN/RP), was modulated by reward feedback but not sensory error feedback during motor adaptation (Palidis et al., 2019). This is consistent with a prominent theory stating that the FRN/RP reflects reward prediction error signals driven by dopamine release (Becker et al., 2014; Carlson et al., 2011; Emeric et al., 2008; Foti et al., 2011; Gehring & Willoughby, 2002; Hauser et al., 2014; Holroyd et al., 2008; Holroyd & Coles, 2002; Mathewson et al., 2008; Miltner et al., 1997; Sambrook & Goslin, 2015, 2016; Vezoli & Procyk, 2009; Walsh & Anderson, 2012; Warren et al., 2015). However, direct evidence for a link between dopamine and the FRN/RP is fairly limited, and no studies have investigated this link in the context of motor adaptation (Enge et al., 2017; Forster et al., 2017; Marco-Pallarés et al., 2009; Mueller et al., 2014; Santesso et al., 2009; Schutte et al., 2020). We hypothesized that levodopa would diminish the magnitude of the FRN/RP along with behavioral expression of reward-based learning in accordance with the “dopamine overdose” hypothesis.

In experiment 2, participants ingested either levodopa or placebo prior to performing a force field adaptation task. We tested for effects of levodopa on savings, in which adaptation is facilitated when a particular perturbation is encountered a second time after washout of initial learning. We also tested for effects of levodopa on anterograde interference, in which adaptation to a force field in a particular direction causes interference with subsequent adaptation to an opposite-direction force field (Bock et al., 2001; Huang et al., 2011; Leow et al., 2012, 2013; R. Christopher Miall et al., 2004; Sing & Smith, 2010). While force field adaptation is thought to rely primarily on sensory error-based learning mechanisms, savings and anterograde interference can be accounted for by additional influences of a reinforcement learning process (Huang et al., 2011). Individuals with Parkinson’s disease show reduced savings and interference despite intact initial adaptation (Bédard & Sanes, 2011; Leow et al., 2012, 2013). While these results suggest a role of dopamine in savings and interference, they typically don’t distinguish between effects of Parkinson’s disease and side-effects of medication. We used pharmacological manipulation in healthy participants to provide a more specific and controlled test for a role of dopamine in savings and interference. We predicted that levodopa would impair savings and interference while leaving initial adaptation unaffected.

We tested for effects of levodopa using a comprehensive battery of motor adaptation tasks. This allowed us to test the hypotheses that dopaminergic mechanisms specifically underlie adaptive motor responses to reward outcomes as well as the formation of motor memories that produce savings and interference effects. We also measured the FRN/RP, a common neural correlate of reward prediction error. This allowed us to test the hypothesis that dopaminergic signaling of reward prediction error in the medial frontal cortex drives reward-based motor adaptation.

## Methods

### Experiment 1

#### Participants

A total of *n=21* [12 female, Age: 20.99 years (SD 3.26)] healthy, right-handed participants were included in experiment 1. All participants were screened for neurological and psychiatric illness, history of drug or alcohol abuse, and contraindications for levodopa. Two participants were excluded due to malfunction of the robot that prevented the experiment from being completed, and two participants were excluded who did not return for the second testing session. Participants provided written informed consent to experimental procedures approved by the Research Ethics Board at Western University.

#### Experimental design

##### Drug administration

All participants underwent two experimental sessions, with levodopa and placebo being administered in separate sessions using a randomized, double-blind, crossover design. The two sessions were separated by a washout period of at least one week. In one session, a capsule was ingested that contained 100 mg of levodopa (L-3,4-dihydroxyphenylalanine) and 25 mg of carbidopa. Levodopa is a dopamine precursor, and carbidopa is a decarboxylase inhibitor given to reduce conversion of levodopa to dopamine in the periphery. This dose has been shown to produce various behavioral effects in healthy young adults (Flöel et al., 2005; Knecht et al., 2004; Onur et al., 2011; Vo et al., 2016, 2017, 2018). In the other session, an equal volume of placebo was administered in an identical capsule. The order of administration was counterbalanced. After administration of the capsule, the robot was calibrated, the EEG cap was placed on the participant’s head, and participants performed a practice block of the behavioral task (see below). Subsequently, the experimental tasks began 45 minutes after ingestion of the capsule to coincide with peak plasma levels of levodopa (Olanow et al., 2000). We measured heart rate, blood pressure, and subjective alertness immediately prior to ingestion of placebo or levodopa and again at the end of each session. Alertness was assessed using the Bond-Lader visual analog scale (Bond & Lader, 1974).

##### Overview of behavioral tasks

Each participant underwent the same experimental tasks in both sessions. Participants made reaching movements toward a visual target and received visual feedback pertaining to reach angle only at movement end point (figure 1). Neural responses to feedback were recorded using EEG. Participants were instructed that each reach terminating within the target would be rewarded with a small monetary bonus. Participants first performed a block of 50 practice trials. The subsequent behavioral procedure consisted of two blocks of a reward learning task and two blocks of a visuomotor rotation (VMR) task. The order of the blocks alternated between the two task types but was otherwise randomized. Participants took self-paced rests between blocks.

**Figure 1.**
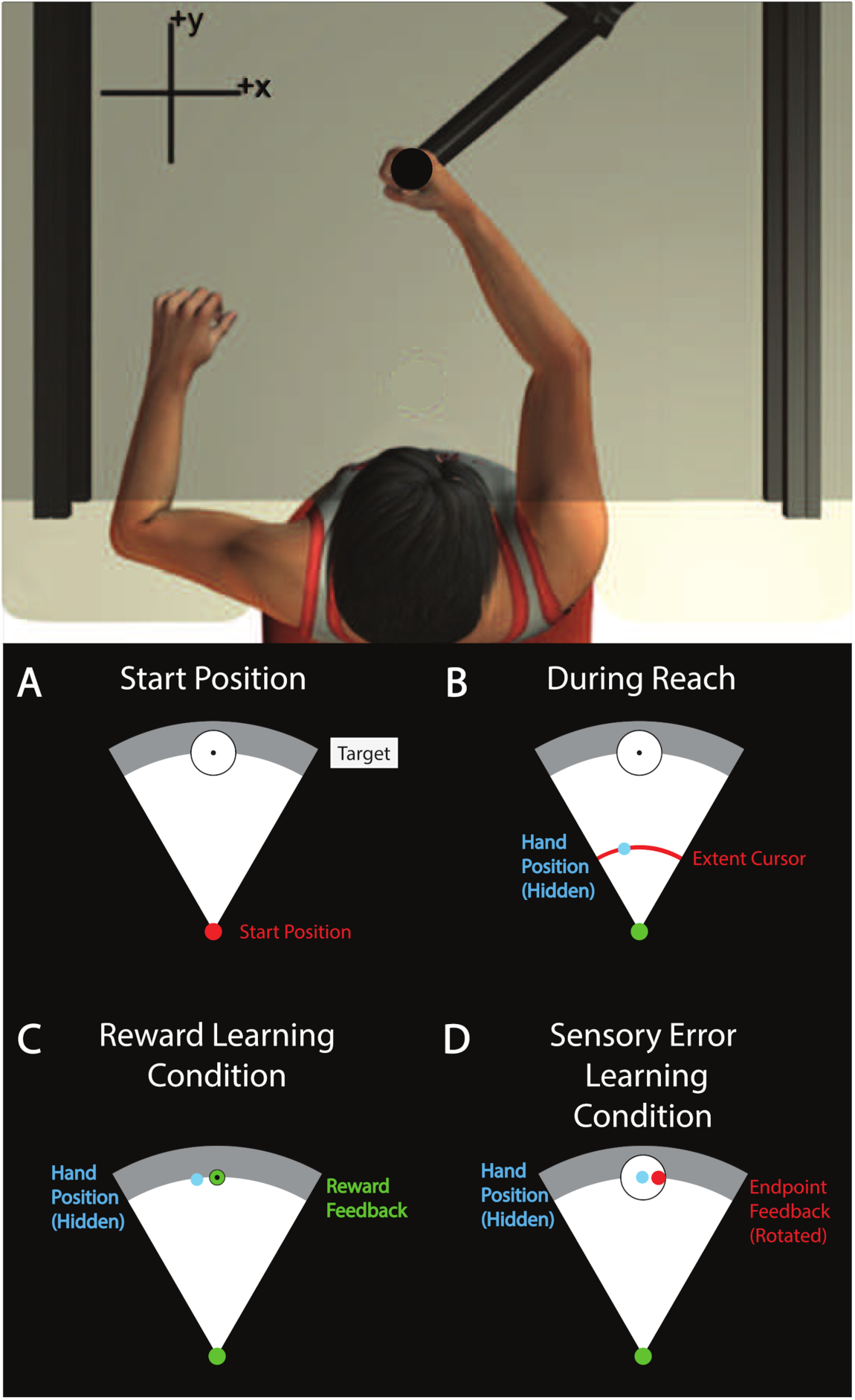
Experimental setup. Top: Apparatus used in both experiments. Participants reached to visual targets while holding the handle of a robotic arm. Vision of the arm was obscured by a screen that displayed visual information related to the task. Bottom: Illustrations of visual display in experiment 1. **A**, Participants made outward reaching movements from a start position at body midline to a visual target. **B**, During reaches, hand position was hidden but an arc-shaped cursor indicated the extent of the reach without revealing reach angle. Feedback was provided at reach end point. ***C***, In the reward learning task, binary feedback represented whether reaches were successful or unsuccessful in hitting the target by turning green or red, respectively. Reach adaptation was induced by providing reward for movements that did not necessarily correspond to the visual target. ***D***, In the visuomotor rotation task, cursor feedback represented the end-point position of the hand. Adaptation was induced by shifting feedback relative to the actual reach angle by rotating it about the start position.

In the VMR task, a cursor appeared at movement end point to represent the position of the hand (Figure 1d). In unperturbed trials, the cursor was displayed directly over the occluded robot handle. In randomly selected trials, the cursor’s position was decoupled from the robot handle position such that the cursor indicated a reach endpoint position that was rotated (about the start position) relative to the actual reach endpoint position. This was intended to produce sensory prediction error and trial-by-trial compensatory changes in reach angle opposite the direction of the rotations. The rotations were small relative to the size of the target, such that participants nearly always landed in the target, fulfilling the goal of the task and earning a monetary reward (the cursor feedback was within the target on 95.5% of trials, SD: 2%). Thus, reward and task error were constant between perturbed and unperturbed feedback, and by comparing the two conditions we could isolate the neural correlates of sensory error processing.

In the reward learning task, no cursor appeared to indicate the position of the hand. Instead, binary feedback represented whether or not participants succeeded in hitting the target (Figure 1c). This allowed us to assess reward-based learning in isolation from sensory error processing, as visual information revealing the position of the hand was not provided. In separate blocks, reward feedback was tailored to produce adaptation towards increasingly clockwise and counterclockwise reach angles. Reward was delivered when the difference between the current reach angle and the median of the previous 10 reach angles was in the direction of intended learning. We compared the neural responses to reward and non-reward feedback to assess the neural correlates of reward processing during adaptation.

#### Apparatus/Behavioral Task

Participants produced reaching movements with their right arm while holding the handle of a robotic arm (InMotion2; Interactive Motion Technologies; figure 1). Position of the robot handle was sampled at 600 Hz. A semi-silvered mirror obscured vision of the arm and displayed visual information related to the task. An air sled supported each participant’s right arm. Participants reached towards a white circular target 14 cm away from a circular start position in front of their chest. The start position turned from red to green to cue the onset of each reach once the handle had remained inside it continuously for 750 ms. Participants were instructed that they must wait for the cue to begin each reach but that it was not necessary to react quickly upon seeing the cue. Participants were instructed to make forward reaches and to stop their hand within the target. An arc-shaped cursor indicated reach extent throughout each movement without revealing reach angle. In only the first five baseline trials of each block, an additional circular cursor continuously indicated the position of the hand throughout the reach. A viscous force field assisted participants in braking their hand when the reach extent was 14 cm. The robot ended each movement by fixing the handle position when the hand velocity decreased below 0.03 m/s. The hand was fixed in place for 700 ms, during which time visual feedback of reach angle was provided. Feedback indicated either reach end point position, a binary reward outcome, or feedback of movement speed (see below). Visual feedback was then removed, and the robot guided the hand back to the start position. Reach end point was defined as the position at which the reach path intersected the perimeter of a circle (14-cm radius) centered at the start position. Reach angle was calculated as the angle between vectors defined by reach end point and the center of the target, each relative to the start position, such that reaching straight ahead corresponds to 0° and counterclockwise reach angles are positive.

Feedback about reach angle was provided either in the form of end-point position feedback or binary reward feedback. The type of feedback, as well as various feedback manipulations, varied according to the assigned experimental block type (see Reward Learning Task and Visuomotor Rotation Task). Participants were told that they would earn additional monetary compensation for reaches that ended within the target, up to a maximum of $10 CAD. Movement duration was defined as the time elapsed between the hand leaving the start position and the moment hand velocity dropped below 0.03 m/s. If movement duration was >700 ms or <450 ms, no feedback pertaining to movement angle was provided. Instead, a gray arc behind the target turned blue or yellow to indicate that the reach was too slow or too fast, respectively. Participants were informed that movements with an incorrect speed would be repeated but would not otherwise affect the experiment. To minimize the impact of eyeblink-related EEG artifacts, participants were asked to fixate their gaze on a black circular target in the center of the reach target and to refrain from blinking throughout each arm movement and subsequent presentation of feedback.

##### Practice block

Each participant first completed a block of practice trials that continued until they achieved 50 movements within the desired range of movement duration. Continuous position feedback was provided during the first 5 trials, and only end-point position feedback was provided for the following 10 trials. Subsequently, no position feedback was provided outside the start position.

##### Reward Learning task

Binary reward feedback was provided to induce adaptation of reach angle (figure 1c). Each session included two blocks in the reward learning condition. The direction of intended learning was clockwise in one block and counterclockwise in the other. Each block continued until the participant completed 125 reaches with acceptable movement duration. Participants reached toward a circular target 1.2 cm in diameter. The first 11 reaches were baseline trials during which continuous position feedback was provided during the first 5 trials, followed by 6 trials with only end-point cursor feedback. After these baseline trials no cursor feedback was provided, and binary reward feedback was instead provided at the end of the movement. Target hits and misses were indicated by the target turning green and red, respectively. Unbeknownst to participants, reward feedback did not necessarily correspond to the visual target. Instead, reward was delivered if the difference between the current reach angle and the median angle of the previous 10 reaches was in the direction of intended learning. When the running median was at least 6° away from zero in the direction of intended learning, reward was delivered at a fixed probability of 50%. This was intended to minimize conscious awareness of the manipulation by limiting adaptation to 6°. Reward was never delivered when the absolute value of the reach angle was greater than 10°, for the same reason. We employed this adaptive, closed-loop reward schedule so that the overall frequency of reward was controlled.

##### Visuomotor rotation task

End-point feedback was rotated relative to the actual reach angle to induce sensory error-based adaptation (figure 1d). Each session included two blocks in the VMR condition. Each block continued until participants completed 124 reaches within acceptable movement duration limits. Participants reached toward a circular target 3.5 cm in diameter. Participants first performed baseline reaches during which cursor feedback reflected veridical reach angle continuously for the first 5 trials and only at movement end point for the subsequent 5 trials. After the baseline reaches the adaptation portion of each block began, unannounced to participants. During the adaptation trials, end-point position feedback was provided indicating a reach angle that was rotated relative to the true reach angle. There were 114 total adaptation trials (38 with 0° rotation, and 19 each with ±2° and ±4° rotations). Participants were instructed that end-point feedback within the target would earn them bonus compensation, but no explicit reward feedback was provided.

#### EEG data acquisition

EEG data were acquired from 16 cap-mounted electrodes with an active electrode system (g.GAMMA; g.tec Medical Engineering) and amplifier (g.USBamp; g.tec Medical Engineering). We recorded from electrodes placed according to the 10-20 System at sites Fp1, Fp2, F3, F4, F7, F8, FT9, FT10, FCz, Cz, C3, C4, CPz, CP3, CP4, and Pz referenced to an electrode placed on participants’ left earlobe. Impedances were maintained below 5 kΩ. Data were sampled at 4,800 Hz and filtered online with band- pass (0.1–1,000 Hz) and notch (60 Hz) filters. A photodiode attached to the display monitor was used to synchronize recordings to stimulus onset.

#### Behavioral data analysis

##### Reward learning task

As in our previous work using a similar task, we computed learning scores in each drug condition by subtracting the average reach angle in the clockwise condition from the average reach angle in the counterclockwise condition (Palidis et al., 2019). As such, positive scores indicate learning. We excluded baseline trials and trials that did not meet the movement duration criteria, as no feedback related to reach angle was provided on these trials. Each block continued until 114 trials after the baseline period met the movement duration criteria, so equal numbers of trials were analyzed for each participant. We tested for the presence of learning by submitting learning scores to 1-sample T-Tests against zero, and we compared learning scores in the placebo and levodopa conditions using paired T-Tests.

We also analyzed trial-by-trial variability in reach angle in response to reinforcement feedback using an approach similar to Pekny et al. (2015). First, we calculated trial-by-trial changes in reach angle as in *Eq. 1*:

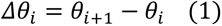

We then multiplied *Δθ_i_* by -1 for trials in the clockwise learning condition, so that positive values for *Δθ_i_* corresponded to changes in reach angle in the direction of intended learning, and any biases in *Δθ* related to the direction of intended learning would have the same sign in the CW and CCW learning conditions. Next, we conditioned *Δθ_i_* on the reinforcement outcome of trial *i* and the drug condition to obtain trial-by-trial changes in reach angle following reward and non-reward after both placebo and levodopa administration. Next, we quantified trial by trial variability in each condition as the natural logarithm of the sample variance of *Δθ_i_*. Our dependent variable is an estimate of variance. This estimate of variance itself has variance due to sampling. For a normal distribution, the variance of a sample variance is proportional to the square of the true population variance. A log transformation is appropriate for linear modeling when the variance of the dependent measure is proportional to the square of its expectation (Montgomery et al., 2021).

We then performed 2x2 repeated measures ANOVA on Log(var(*Δθ_i_*)). The factors were drug (levels: placebo, levodopa), and reward outcome on trial *i* (levels: non-reward, reward).

##### Visuomotor rotation task

To quantify trial-by-trial learning we first calculated the change in reach angle between successive trials, as in *Eq. 1*. We then performed a linear regression on *Δθ_i_* with the rotation imposed on trial *i* as the predictor variable. The rotation was 0°, ±2°, or ±4°. This regression was performed on an individual participant basis, separately for placebo and levodopa conditions. We excluded trials that did not meet the duration criteria as no visual feedback was provided on these trials. We took the resulting slope estimates multiplied by -1 as a metric of learning rate, as it reflects the portion of visual errors that participants corrected with a trial-by-trial adaptive process. We tested for the presence of adaptation in each condition by submitting learning rates to 1-sample t-tests against zero. We tested for an effect of levodopa vs placebo on learning rates using a paired t-test.

#### EEG preprocessing

EEG data were resampled to 480 Hz and filtered off-line between 0.1 and 35 Hz with a second-order Butterworth filter. Continuous data were segmented into 2-s epochs time-locked to feedback stimulus onset at 0 ms (time range: -500 to +1,500 ms). Epochs flagged for containing artifacts as well as any channels with bad recordings were removed after visual inspection. One participant was excluded entirely from the EEG analysis due to excessive muscle artifacts. Subsequently, extended infomax independent component analysis was performed on each participant’s data (Delorme & Makeig, 2004). Components reflecting eye movements and blink artifacts were identified by visual inspection and subtracted by projection of the remaining components back to the voltage time series.

#### EEG data analysis

After artifact removal, we computed ERPs by trial averaging EEG time series epochs for various feedback conditions described in the sections below. ERPs were computed on an individual participant basis separately for recordings from channels FCz and Pz. We selected FCz and Pz a priori because these electrodes typically correspond to the peaks of the scalp distributions for the feedback related negativity/reward positivity and the P300 ERP components, respectively. We found this to be true in a previous experiment using a very similar paradigm (Palidis et al., 2019). All ERPs were baseline corrected by subtracting the average voltage in the 75-ms period immediately following stimulus onset. We used a baseline period following stimulus onset because stimuli were presented immediately upon movement termination and the period before stimulus presentation was more likely to be affected by movement related artifacts. Trials in which reaches did not meet the movement duration criteria were excluded, as feedback relevant to reach adaptation was not provided on these trials. Finally, ERPs were low-pass filtered with a cutoff frequency of 30 Hz.

We computed ERPs separately following administration of placebo and levodopa. In the reward learning task, we computed ERPs separately for feedback indicating non-reward (placebo: 107.2 ±9.7 trials, levodopa: 104.0 ±8.3 trials) and feedback indicating reward (placebo: 118.4 ±9.6 trials, levodopa: 118.0 ±8.1 trials). In the visuomotor rotation task, we computed ERPs separately for veridical endpoint feedback (placebo: 72.6 ± 3.5 trials, levodopa: 72.9 ± 3.1 trials), ±2° rotated feedback (placebo: 70.8 ± 5.2 trials, levodopa: 72.1 ± 3.8 trials), and ±4° rotated feedback (placebo: 64.5 ± 4.7 trials, levodopa: 66.3 ± 4.1 trials). We excluded trials in which the cursor did not land within the target.

We selected time windows of interest for ERP analysis using independent data from a previous experiment with very similar procedures (Palidis et al., 2019). We analyzed the amplitudes of FRN/RP and P300 components within 50 ms time windows centered around the latencies of the FRN/RP and P300 peaks observed in our previous study. The FNR/RP peak was taken as the maximum value of the difference between ERPs elicited by reward and non-reward feedback recorded from electrode FCz (latency: 292ms). For completeness, we used the same time window to test for FRN/RP effects in the visuomotor rotation task of the current study although we did not observe an FRN/RP component in our previous visuomotor rotation task. The P300 peak latencies were determined separately for reward and non-reward feedback as the times of maximal amplitude of ERPs recorded from electrode Pz (reward: 319ms, non-reward: 371ms). The peak latencies selected for the FRN/RP and P300 components in the reward learning task corresponded very closely to the peaks observed in the current data. However, the P300 peak in the visuomotor rotation task of the current study was earlier than that in our previous experiment. This difference in latency may be due to changes in the nature of the feedback. Thus, we determined the latency of the P300 peak in the visuomotor rotation task of the current study using a data-driven method that does not bias comparisons between conditions (Brooks et al., 2017). We aggregated all trials across conditions and participants and computed a trial averaged ERP using recordings from electrode Pz. The P300 peak was determined as the maximal amplitude of this averaged waveform (latency: 317ms). This method is only suitable for comparing waveforms of different amplitude but similar morphology across conditions, and thus could not be applied to the ERPs in the reward learning task (Brooks et al., 2017).

We tested for effects of feedback manipulations on FRN/RP components using the average amplitude of ERPs recorded from electrode FCz within the FRN/RP time window. We tested for effects on P300 ERP components using average amplitude of ERPs recorded from electrode Pz within the P300 time window corresponding to a given condition. For the reward learning task, we used 2x2 repeated measures ANOVAs with factors drug (levels: placebo, levodopa) and reinforcement outcome (levels: reward, non-reward). For the visuomotor rotation task, we used 2x3 repeated measures ANOVAs with factors drug (levels: placebo, Levodopa), and rotation (levels: 0°, ±2°, ±4°).

### Experiment 2

#### Participants

A total of 38 participants were included in experiment 2 (Table 2). All participants were screened for neurological and psychiatric illness, history of drug or alcohol abuse, and contraindications for levodopa. Participants provided written informed consent to experimental procedures approved by the Research Ethics Board at Western University.

#### Procedure

##### Drug administration

Participants were administered either levodopa or placebo in a randomized double-blind design. A capsule was ingested that contained 100 mg of levodopa (L-3,4-dihydroxyphenylalanine) and 25 mg of carbidopa or an equal volume of placebo. The experimental tasks began 45 minutes after ingestion of the capsule to coincide with peak plasma levels of levodopa. We measured subjective alertness using the Bond-Lader visual analog scale (Bond & Lader, 1974) as well as heart rate and blood pressure immediately prior to ingesting the capsule and again at the end of each session.

##### Force field adaptation task

Participants produced reaching movements with their right arm while holding the handle of a robotic arm (InMotion2; Interactive Motion Technologies). The position of the robot handle was sampled at 600 Hz. A semi-silvered mirror obscured vision of the arm and displayed visual information related to the task. An air sled supported each participant’s right arm.

On each trial, participants reached from a central home position (blue circle 20 mm in diameter) to one of 8 circular targets (24 mm in diameter) arranged around the home position at a distance of 10 cm. The target angles were 0°, 45°, 90°, 135°, 180°, 225°, 270°, and 315°. A 5-mm pink circular cursor represented the position of the robot handle. When the cursor reached the target on each trial, the target either turned blue to indicate that the movement duration was satisfactory (375 ± 100 ms), green to indicate that the movement was too slow, or red to indicate that the movement was too fast. The subject moved the robot handle back to the home position at the end of each reach.

In null field blocks, the robot motors did not apply any external forces to the hand. In force field blocks, the robot applied forces to the hand that were perpendicular to the direction of movement and proportional to the velocity of the hand (*eq. 2)*. The direction of the force field was either clockwise or counterclockwise, in separate blocks.

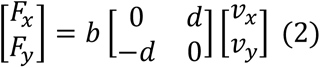

*x* and *y* correspond to the lateral and sagittal directions. *F_x_* and *F_y_* describe the forces applied to the hand, *v_x_* and *v_y_* describe the velocity of the hand, *b* is the field constant, and *d* corresponds to the direction (*d* = 1 for a clockwise force field (CWFF), -1 for a counterclockwise force field (CCWFF) or 0 for a null field (NF)).

All participants completed five blocks of 96 trials. Each block consisted of 12 reaches to each of the 8 targets presented in random order. The five blocks occurred in the following order: NFa (null field), FF1a (CWFF), NFb (null field), FF1b (CWFF), FF2 (CCWFF). Trials 6, 24, 35, 50, 71, and 91 of each block were “catch trials”, during which reaches occurred in a null field. When the force field is suddenly removed in catch trials, errors occur in the opposite direction of the force field. A reduction in reach error during force field trials may reflect either adaptation to the force field, stiffening of the arm, or changes in feedback corrections. The magnitude of errors opposite the force field in catch trials is thought to better capture adaptation of feedforward control. Similar to catch trials, we expected after-effects at the beginning of NFa in the form of counterclockwise reach errors after the sudden removal of the clockwise force field in FF1a.

#### Data analysis

Robot handle positional data were low-pass filtered with a 40 Hz cutoff frequency and differentiated to yield instantaneous velocity and acceleration. On each trial, movement onset and end of movement were defined according to a velocity threshold set at 5% of the maximum tangential velocity of the robot endpoint. Our behavioral measure of interest was the lateral deviation of the hand at the time of peak tangential velocity. Perpendicular deviation (PD) was calculated relative to a line drawn from the position of movement onset in the direction of the target angle (either 0°, 45°, 90°, 135°, 180°, 225°, 270°, or 315°). PD was calculated for each trial as the perpendicular distance between the position of the hand at peak velocity and this line, with positive PD corresponding to clockwise deviations. For non-catch trials, PD was averaged across trials within 12 bins of 8 trials each. We analyzed effects related to adaptation separately for an early and late period of each block. The early period consisted of the first 5 bins (trials 1-40, catch trials: 6,24,35) and the late period consisted of the remaining 7 bins (trials 41-96, catch trials: 50,71,91). Baseline PD was computed as the average PD in the late period of NFa. We computed metrics for adaptation, savings, after-effects, and learning with interference separately for the early and late periods, and separately for catch trials and non-catch trials. All metrics were computed so that positive values corresponded to the effects of interest, and values of zero correspond to no effect. We tested for adaptation, savings, after-effects, and learning with interference using 1-sample t-tests against zero. We tested for differences between the placebo and levodopa groups using paired t-tests.

##### Non-catch trials

Adaptation metrics were computed to capture reductions in error during FF1a relative to the initial errors caused by the onset of the force field. Our measure of early adaptation was the average PD in the first bin of FF1a minus the average PD across subsequent bins within the early period of FF1a (bins 2-5). Our measure of late adaptation was the average PD in the first bin of FF1a minus the average PD across bins in the late period of FF1a (bins 6-12). Savings metrics were computed to measure reductions in errors during the second exposure to FF1 compared to the first. Savings was measured as the difference in PD between FF1a and FF1b (FF1a – FF1b), separately for PD averaged across bins within the early and late periods. Adaptation to FF1a caused after-effects in the form of errors upon its sudden removal at the onset of NFb. After-effects were measured as the difference between baseline PD and the PD in NFb (baseline – NFb), separately for PD averaged across bins in the early and late periods of NFb. We expected large initial errors at the onset of FF2 due to a combination of after-effects from the removal of FF1b and the introduction of a novel force field. Previous adaptation to FF1b was also expected to cause anterograde interference during adaptation to FF2 as the force fields were opposite. Metrics for adaptation with interference were computed to capture reductions in errors during FF2 relative to the initial errors caused by the onset of the force field. Early adaptation with interference was measured by subtracting the average PD from the first bin of FF2 from the average PD across subsequent bins within the early period of FF2 (bins 2-5). Late adaptation with interference was measured by subtracting the average PD in the first bin of FF2 from the average PD across subsequent bins in the late period of FF2 (bins 6-12).

##### Catch trials

When a force field is suddenly removed during catch trials, adaptation to the force field is reflected in errors opposite the direction of the force field. Adaptation effects were computed as the baseline PD minus the PD in FF1a averaged across catch trials, separately for catch trials in the early and late period. Improved adaptation due to savings was expected to cause larger errors in catch trial during FF1b compared to FF1a. Savings was computed as the PD in FF1a minus the PD in FF1b, averaged across catch trials separately for the early and late periods. Learning effects with interference were computed using data from FF2. There was no suitable baseline PD to analyze learning in this block. Instead, the PD of the first catch trial was subtracted from the PD of each of the later catch trials, separately for catch trials in the early and late periods. This captures changes in catch trial PD opposite the direction of FF2 due to adaptation.

#### Statistics

Statistical tests were implemented using JASP v0.14.1. We compared sample means using 1 sample T-Tests, paired sample T-Tests, or independent sample T-Tests. These comparisons allowed us to compute one-tailed Bayes factors representing *p*;(*data*|*H*_+_) / *p*;(*data*|*H*_0_), where *H*_0_ represents the null hypothesis corresponding to the standard *t*-distribution for an effect size of 0, and *H*_+_ represents the alternative hypothesis corresponding to a *t*-distribution constructed using a one-tailed prior distribution of effect sizes. The use of 1-tailed priors is recommended in the case of directional hypotheses to provide “a fairer balance between the ability to provide evidence for *H*0 and *H*1” (Keysers et al., 2020). We used the default effect size priors implemented in JASP (Cauchy scale 0.707). These priors are generally appropriate for effect sizes typical of neuroscience research, and the use of default priors is recommended for standardized and objective analysis (Keysers et al., 2020; Rouder et al., 2012; Wetzels et al., 2011). Bayesian estimates of effect size are reported as median posterior Cohen’s δ with 95% credibility interval using 2-tailed priors for H1 to avoid biasing the estimate in the expected direction. We also report T-statistics, p-values, and 95% confidence intervals generated using 2-tailed frequentist T-Tests. For factorial analyses, we conducted frequentist and Bayesian repeated measures ANOVAs using JASP with default priors. Bayes factors were computed for the inclusion of each effect as the ratio of the data likelihood under the model containing that effect vs equivalent models stripped of that effect. Bayes factors >3 and >10 were taken as moderate and strong evidence in favor of the alternative hypothesis, respectively. Bayes factors <1/3 and <1/10 were taken as moderate and strong evidence in favor of the null hypothesis, respectively. Bayes factors between 1/3 and 3 were taken as inconclusive evidence (Keysers et al., 2020).

Directional priors used for alternative hypotheses specified our predictions that learning metrics would be greater than zero (Reward learning score, VMR learning rate, force field adaptation, savings, after-effects, and adaptation with interference). In comparing placebo and levodopa conditions, our alternative hypotheses specified that learning metrics would be lower in levodopa conditions than placebo conditions, in accordance with the “dopamine overdose” hypothesis. The only exception was that we predicted adaptation with interference would be increased by levodopa. If anterograde interference is caused by dopaminergic reinforcement learning, then the “dopamine overdose” effect should reduce interference and facilitate adaptation. All other Bayes factors are computed with 2-tailed priors, as they were conducted without directional a priori hypotheses (control measures, etc.).

## Results

### Experiment 1

#### Control measures

Participants’ judgments at the end of the second session as to whether they received placebo or drug were correct at near chance level (47.62%). Table 1 shows the values for heart rate, blood pressure, and alertness recorded at the beginning and end of each experimental session for both the placebo and levodopa conditions. We computed the percent change in heart rate and blood pressure recorded at the beginning and end of each session. There were no reliable differences between the levodopa and placebo conditions in the percent change of heart rate (t(18) = 0.70, p=0.49, 95%CI for difference = [-0.03 0.07], BF = 0.30, posterior δ: median = 0.139, 95%CI = [-0.278 0.565]), systolic blood pressure (t(18) = -0.39, p=0.70, 95%CI for difference = [-0.06 0.04], BF = 0.25, posterior δ: median = -0.077, 95%CI = [-0.498 0.338]), or diastolic blood pressure (t(18) = -0.88, p=0.39, 95%CI for difference = [-0.07 0.03], BF = 0.33, posterior δ: median = -0.173, 95%CI = [-0.603 0.245]). We did observe a significant difference between levodopa and placebo in the percent change of alertness (t(20) = 2.46, p=0.023, 95%CI for difference = [0.02 0.19], BF = 2.53, posterior δ: median = 0.477, 95%CI = [0.044 0.930]). However, this effect was likely due to chance as alertness was only different between the two drug conditions at the time point pre-administration of the capsule (t(20) = 2.18, p=0.042), but not post-administration (t(20) = -0.068, p=0.95). We also tested for effects of levodopa on the median response time (the latency between the go cue and the robot handle leaving the home position), and the median movement time (table 1). We observed no reliable differences in response time between the placebo and levodopa conditions in either the reward learning task (t(20)=0.72, p=0.48, 95%CI for difference = [-37.49 77.34], BF = 0.29, posterior δ: median = 0.137, 95%CI = [-0.261 0.545]), or the VMR task (t(20)=0.62, p=0.54, 95%CI for difference = [-33.91 62.56], BF = 0.27, posterior δ: median = 0.118, 95%CI = [-0.280 0.523]). We also observed no reliable difference in movement time between the placebo and levodopa conditions in either the reward learning task (t(20)=- 0.11, p=0.91, 95%CI for difference = [-20.75 18.69], BF = 0.23, posterior δ: median = - 0.021, 95%CI = [-0.420 0.377]), or the VMR task (t(20)=-0.21, p=0.84, 95%CI for difference = [-16.21 13.27], BF = 0.23, posterior δ: median = -0.039, 95%CI = [-0.44 0.358]).

**Table 1:** Control measurements from experiment 1. Heart rate (bpm). Systolic blood pressure (mm Hg). Diastolic blood pressure (mm Hg). Alertness, Bond-Lader visual analog scale alertness measure. Response Time, latency between go cue and hand exiting the start position (ms). Movement Time, duration of movement (ms). RL, reward learning task. VMR, visuomotor rotation task.

| Measure | Placebo | Levodopa |
| --- | --- | --- |
| Heart Rate | Pre: 76.24 (SD: 11.29)<br>Post: 69.60 (SD: 7.27) | Pre: 77.55 (SD: 8.41)<br>Post: 71.53 (SD: 6.92) |
| Systolic | Pre: 104.43 (SD: 9.01)<br>Post: 104.20 (SD: 6.47) | Pre: 103.95 (SD: 8.34)<br>Post: 102.79 (SD: 8.70) |
| Diastolic | Pre: 72.14 (SD: 5.14)<br>Post: 73.20 (SD: 4.55) | Pre: 70.55 (SD: 6.81)<br>Post: 69.74 (SD: 6.04) |
| Alertness | Pre: 64.58 (SD: 8.38)<br>Post: 47.99 (SD: 15.43) | Pre: 58.20 (SD: 11.79)<br>Post: 48.16 (SD: 15.33) |
| Response Time | RL: 464.09 (SD: 140.05)<br>VMR: 445.91 (SD: 120.96) | RL: 484.01 (SD: 149.00)<br>VMR: 460.24 (SD: 133.04) |
| Movement Time | RL: 548.17 (SD: 37.04)<br>VMR: 547.90 (SD:34.92) | RL: 547.14 (SD: 35.28)<br>VMR: 546.43 (SD: 40.52) |

**Table 2:** Control measurements from Experiment 2. Heart rate (bpm). Systolic blood pressure (mm Hg). Diastolic blood pressure (mm Hg). Alertness, Bond-Lader visual analog scale alertness measure. Peak Velocity, maximum tangential velocity of the hand averaged across trials (m/s).

| Measure | Placebo | Levodopa |
| --- | --- | --- |
| *n* | 19 | 19 |
| *n* female | 9 | 10 |
| Age | 21.2 (SD: 2.5) | 22.2 (SD: 3.4 years) |
| Heart Rate | Pre: 75.1 (SD: 9.5)<br>Post: 66.2 (SD: 10.2) | Pre: 71.6842 (SD: 12.8)<br>Post: 65.7 (SD: 11.3) |
| Systolic | Pre: 109.2 (SD: 15.4)<br>Post: 104.8 (SD: 14.5) | Pre: 108.4 (SD: 11.4)<br>Post: 99.7 (SD: 10.1) |
| Diastolic | Pre: 72.0 (SD: 10.2)<br>Post: 70.1 (SD: 10.2) | Pre: 73.2 (SD: 15.5)<br>Post: 67.0 (SD: 8.2) |
| Alertness | Pre: 31.3 (SD: 15.3)<br>Post: 39.4 (SD: 17.0) | Pre: 27.1 (SD: 11.0)<br>Post: 43.4 (SD: 12.7) |
| Peak Velocity | 0.43 (SD = 0.01) | 0.43 (SD = 0.02) |

### Behavioral results

#### Reward learning task

Behavioral data from the reward learning task are shown in Figure 2. Learning scores were reliably greater than zero in both the placebo condition (mean = 6.03, SD = 3.58, t(20) = 7.72, p = 2.02e-7, 95%CI = [4.40 7.66], BF = 1.56e5, posterior δ: median = 1.58 95%CI = [0.92 2.28]), and the levodopa condition (mean = 6.93, SD = 3.86, t(20) = 8.23, p = 7.49e-8, 95%CI = [5.17 8.69], BF = 3.9e5, posterior δ: median = 1.69, 95%CI = [1.00 2.41]) conditions. Learning scores were slightly higher in the levodopa condition, though this difference was not statistically reliable. This result provided strong evidence against our hypothesis of reduced learning in the levodopa group (t(20) = -1.58, p = 0.13, 95%CI for difference = [-2.09 0.29], BF = 0.10, posterior δ: median = -0.30, 95%CI = [-0.73 0.11]). We observed similar evidence against the hypothesized effect of levodopa when learning scores were computed using only the final 20 trials in each block (t(20) = -1.60, p = 0.13, 95%CI for difference = [-3.05 0.40], BF = 0.10, posterior δ: median = -0.31, 95%CI = [-0.73 0.10]).

**Figure 2.**
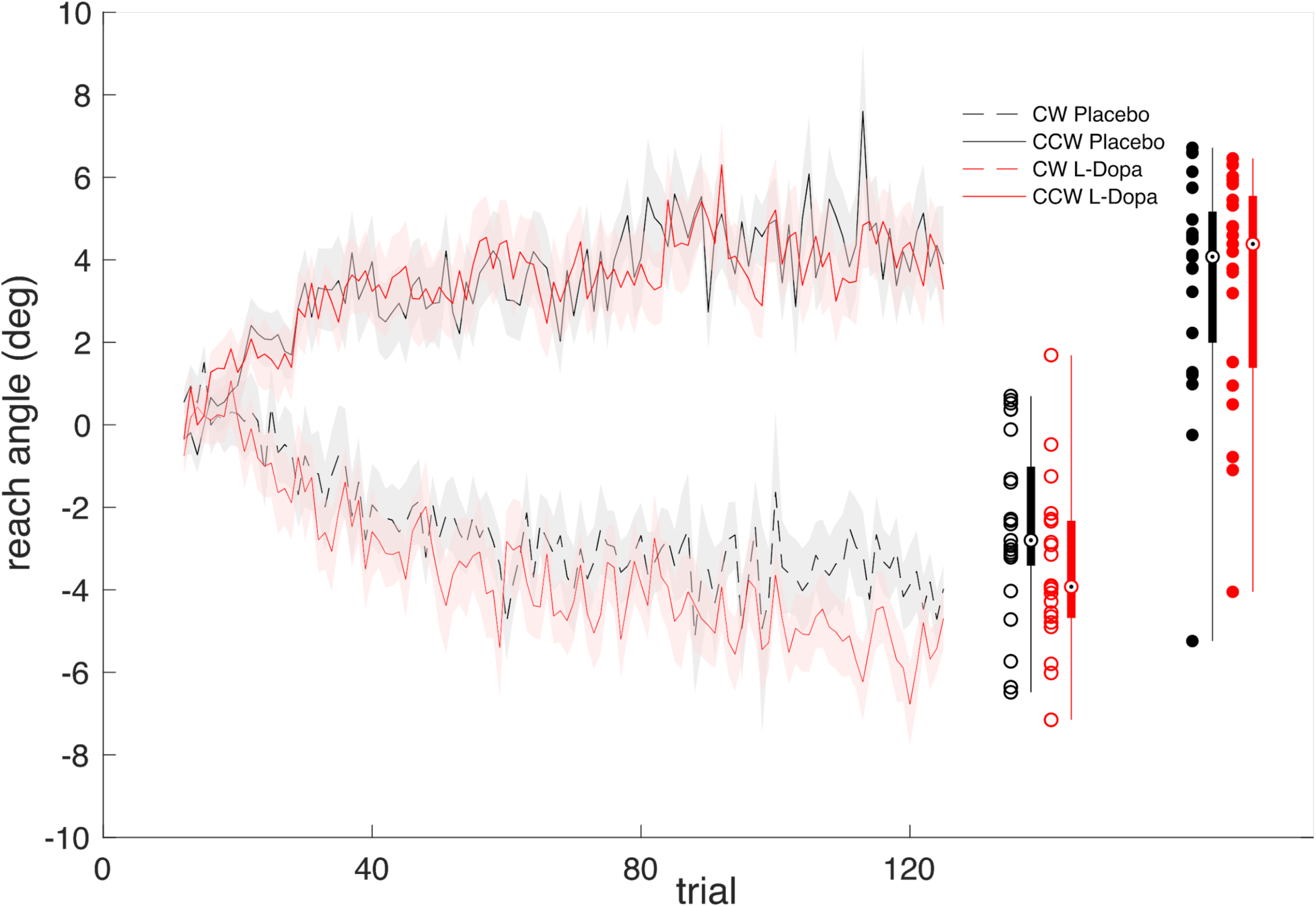
Reward-based motor adaptation (n=21). The time series show group average reach angles in the reward learning task across trials (Shaded region: ± SEM). After both placebo and levodopa administration, participants completed a block in each direction of intended learning condition [clockwise (CW) and counterclockwise (CCW)]. Trials 1-11 were baseline trials without reinforcement feedback, and are not shown. Individual data points on the right show the average reach angles across trials in each condition for each participant (CCW: solid markers, CW: open markers, black: placebo, red: L-Dopa). Box plots summarize the distributions of individual data using circular markers to indicate the medians, thick lines to indicate interquartile ranges, and thin lines to indicate full ranges.

The variability of trial-by-trial changes in reach angle following reward and non-reward outcomes is shown in Figure 3. We found a reliable main effect of reinforcement outcome on the log transformed variance of trial-by-trial changes in reach angle (F(1,20) = 74.84, p = 3.41e-8, 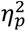= 0.79, BF = 3.02e14). This indicates an increase in trial-by-trial variance of reach angle following non-reward outcomes relative to reward. We found moderate evidence against effects of drug condition (F(1,20) = 0.0072, p = 0.93, 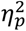= 3.86e-4, BF = 0.22) and reward by drug interaction (F(1,20) = 0.0478, p = 0.829,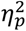=2.38e-3, BF = 0.30).

**Figure 3.**
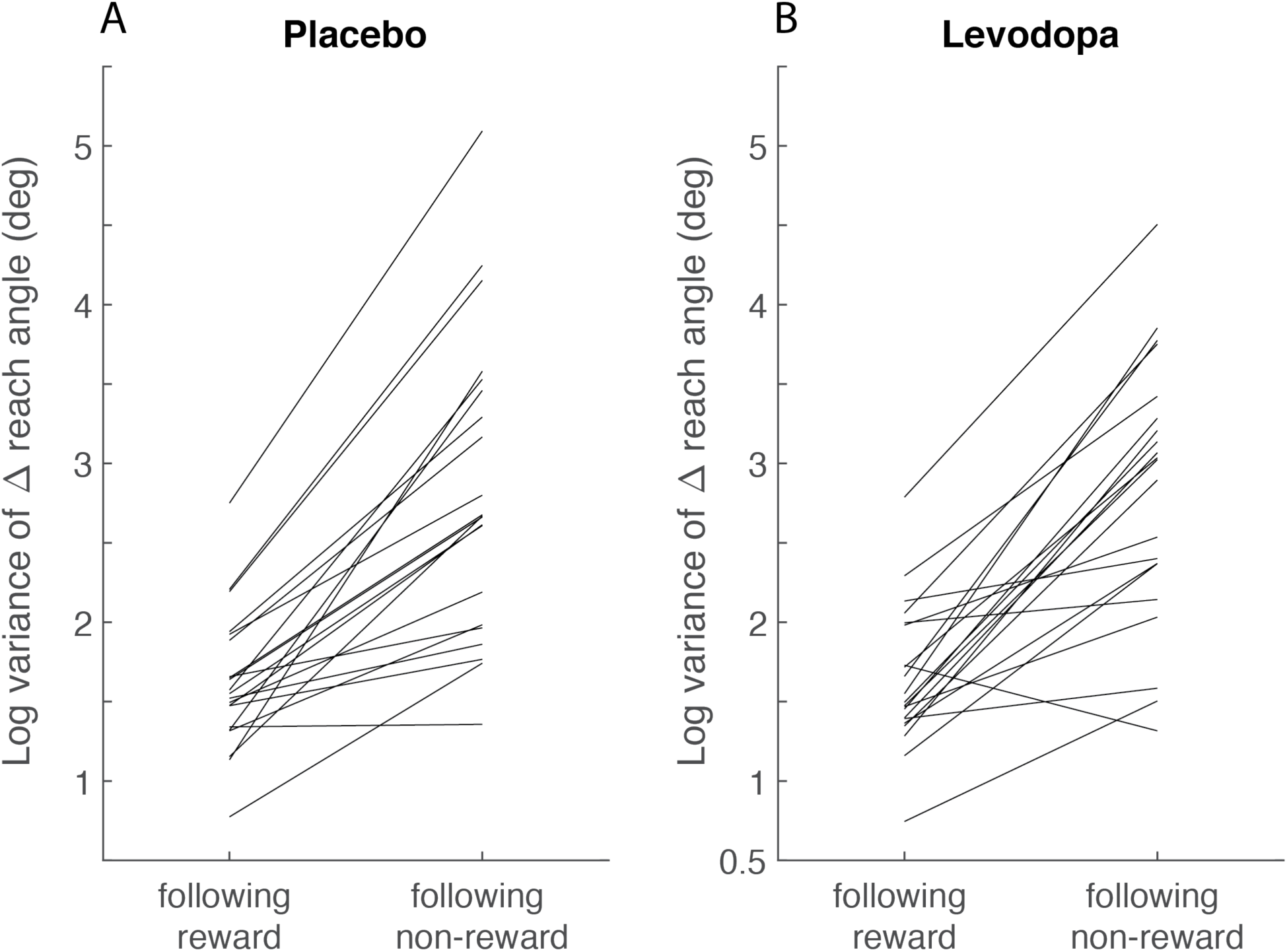
Reward induced modulation of trial-by-trial variability of reach angle (n=21). The log transformed variance of trial-by-trial changes in reach angle (deg) following reward and non-reward are plotted for each participant following administration of levodopa (**A)** and placebo (**B)**.

#### Visuomotor rotation task

Mean trial-by-trial changes in reach angle after the different feedback rotations are shown in Figure 4. Learning rates were reliably greater than zero following administration of both placebo (mean: 0.313, SD: 0.133, t(20) = 10.77, p = 8.93e-10, 95%CI = [0.25 0.37], BF = 2.4e7, posterior δ: median = 2.22, 95%CI = [1.40 3.10])) and levodopa (mean: 0.294, SD: 0.102, t(20) = 13.18, p = 2.54e-11, 95%CI = [0.25 0.34], BF = 6.75e8). Learning rates were not reliably different in the two conditions (t(20) = 0.703, p=0.491, 95%CI for difference = [-0.04 0.07], BF = 0.42, posterior δ: median = 0.134, 95%CI = [-0.265 0.540])).

**Figure 4.**
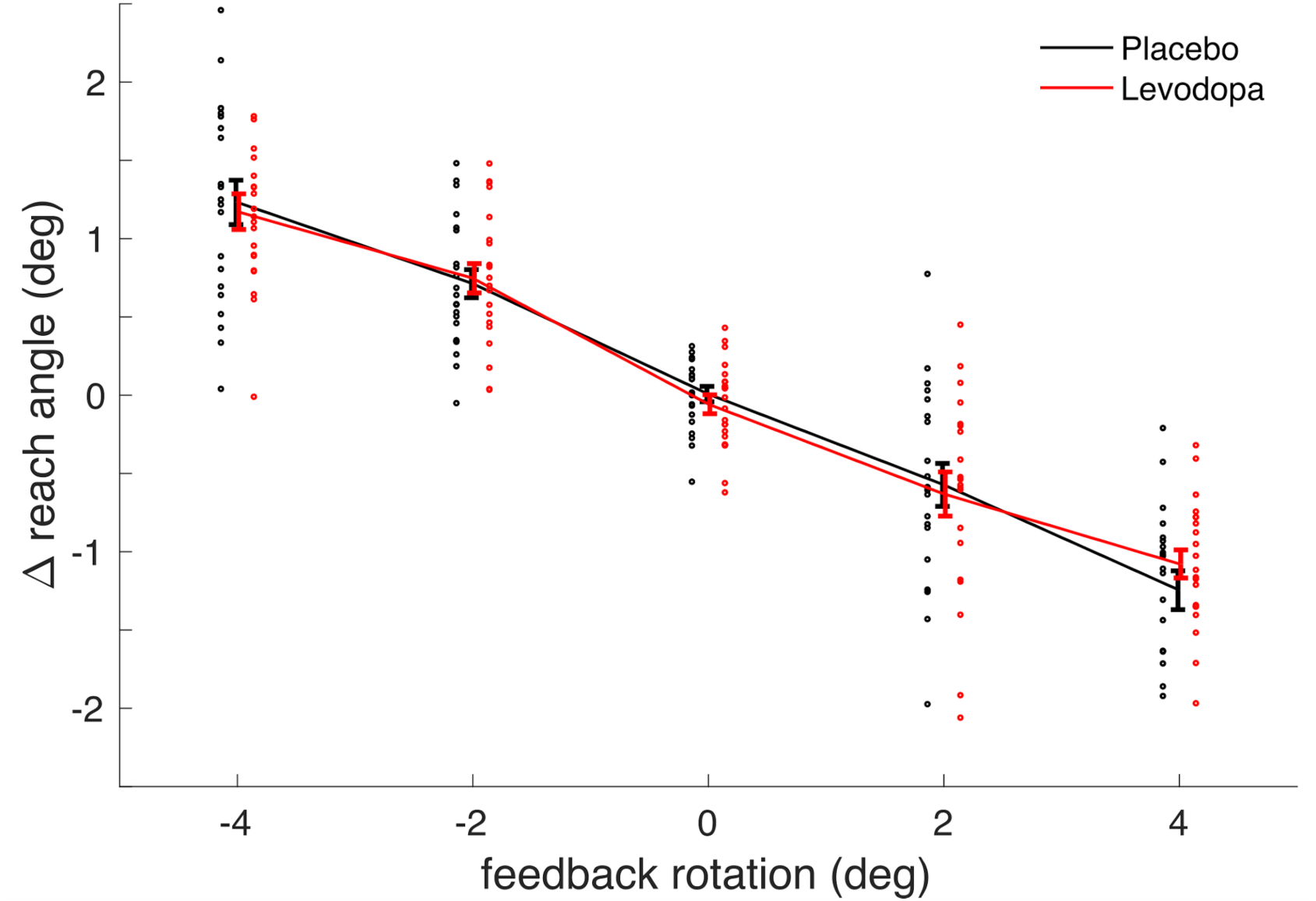
Sensory error-based motor adaptation (n=21). The average change in reach angle between subsequent pairs of trials is plotted for each size and direction of rotation imposed on the preceding trial. The average change in reach angle is in all cases opposite to the rotation, indicating that participants adapted their reaches to counteract the perturbations. Individual data points show average changes in reach angle across trials for each participant. Lines show average change in reach angle across participants (Error bars: ± SEM).

### Event-related potential results

#### Reward learning task.

Feedback-related negativity/Reward positivity: Event-related potentials (ERPs) elicited by reinforcement feedback at electrode FCz are shown in Figure 5a. We analyzed the FRN/RP by submitting the average ERP amplitude at electrode FCz between 267- 317ms to frequentist and bayesian repeated measures ANOVAs (figure 5b). We found a reliable main effect of reward outcome on FRN/RP amplitude (F(1,19) = 42.25, p = 3.16e-6, 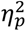=0.69, BF = 8.89e8). We observed moderate evidence both against an effect of drug (F(1,19) = 0.13, p = 0.73, 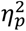=6.56e-3, BF = 0.24) and a reward by drug interaction (F(1,19) = 0.2, p = 0.66, 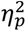 =0.01, BF = 0.30) on FRN/RP amplitude.

**Figure 5.**
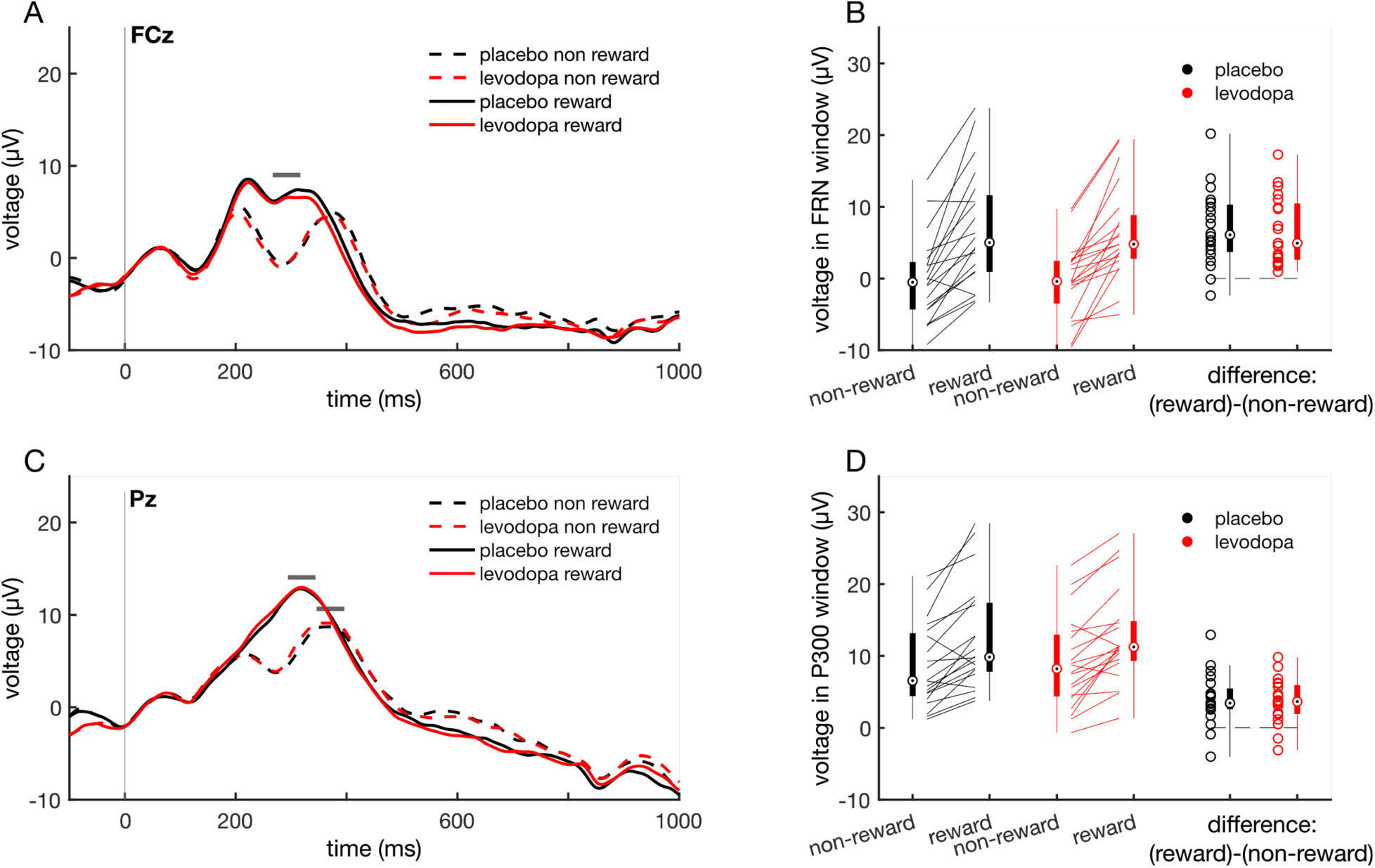
Event-related potentials elicited by reinforcement feedback (n=20). **A**, Grand averaged ERPs recorded from electrode FCz. ERPs are aligned to reinforcement feedback presentation (0 ms: vertical grey line). Horizontal grey bar indicates time window for FRN/RP analysis (267-317ms). Trials were selected by reinforcement outcome (reward or non-reward) and drug condition (levodopa or placebo) for ERP averaging. **B**, ERP amplitude during the FRN/RP time window. Individual participants’ data show amplitude following reward, non-reward, and the difference [(reward) - (non- reward)]. Boxplots indicate the median (circular markers), the interquartile range (thick bars) and the range (thin lines). **C**, Trial averaged ERPs recorded from electrode Pz. Horizontal grey bars indicate time windows for P300 analysis (Reward: 294-344ms, Non-reward: 346-396ms). **D**, ERP amplitudes during the P300 time windows, as in B.

P300: ERPs elicited by reinforcement feedback at electrode Pz are shown in Figure 5c. We analyzed the P300 by submitting the average ERP amplitudes at electrode Pz during the P300 time windows (Reward: 294-344ms, Non-reward: 346-396ms) to frequentist and bayesian repeated measures ANOVAs (figure 5d). We found a reliable main effect of reward outcome on P300 amplitude (F(1,19) = 35.83, p = 9.26e-6, 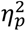=0.65, BF = 3.5e5). We observed moderate evidence both against an effect of drug (F(1,19) = 0.20, p = 0.66, 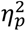=0.01, BF = 0.26) and against a reward by drug interaction (F(1,19) = 0.13, p = 0.73, 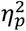= 6.56e-3, BF = 0.29) on P300 amplitudes.

#### Visuomotor rotation task.

Feedback-related negativity/Reward positivity: ERPs elicited by endpoint cursor feedback at electrode FCz are shown in Figure 6a. We analyzed the FRN/RP by submitting the average ERP amplitude at electrode FCz between 267-317ms to repeated measures ANOVAs (figure 6b). We did not find reliable main effects of drug (F(1,19) = 1.37, p = 0.26, 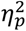=0.07), or feedback rotation (F(2,38) = 0.1, p = 0.86 (Greenhouse-Geisser corrected), 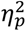= 5.12e-3). We did observe a reliable drug by rotation interaction effect (F(2,38) = 4.75, p = 0.02 (Greenhouse-Geisser corrected), 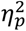=0.2). Simple main effects did not show reliable main effects of rotation in either the placebo (F(2,38) = 2.17, p=0.13) or levodopa (F(2,38) = 2.06, p=0.14) conditions on FRN/RP amplitudes.

**Figure 6.**
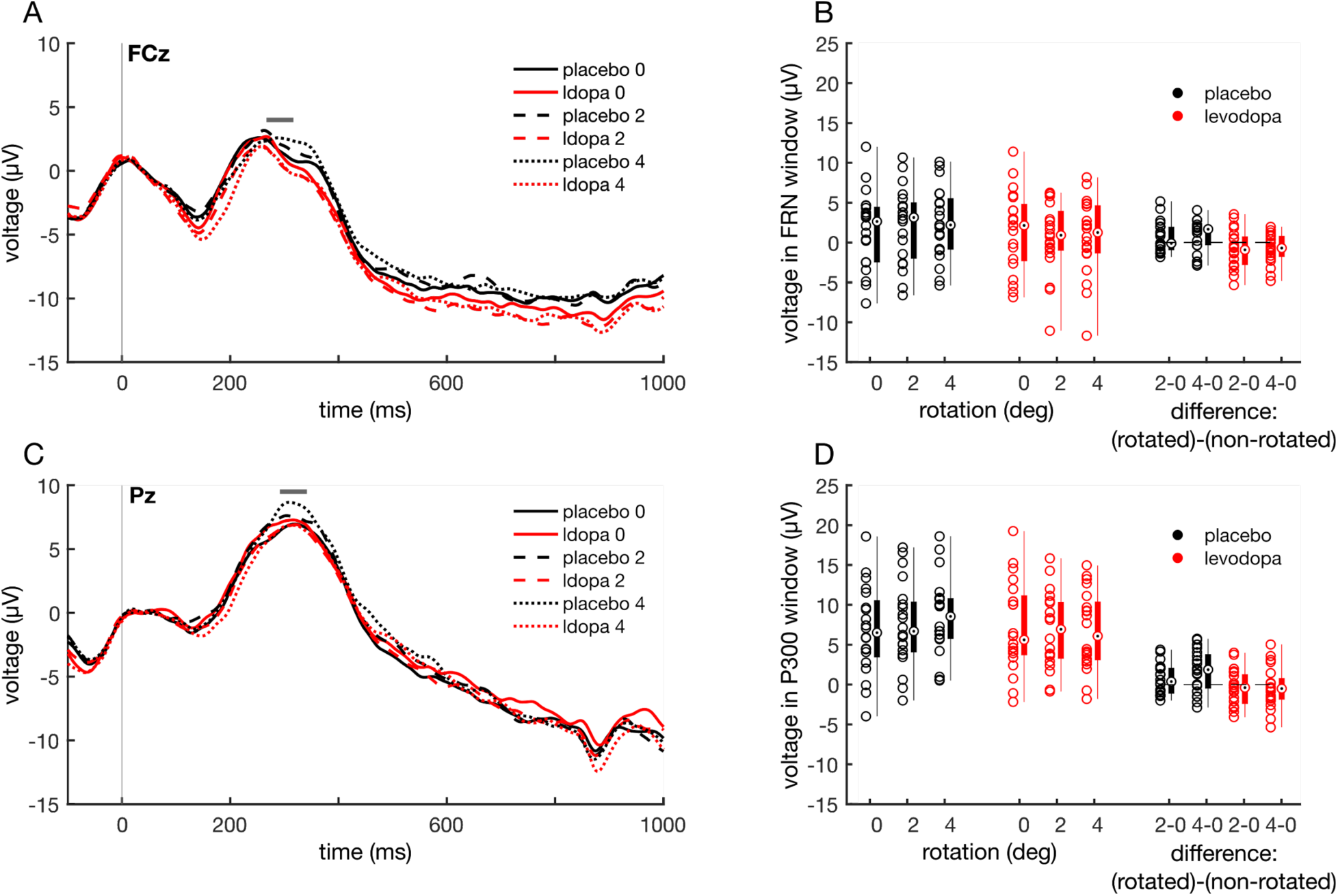
Event-related potentials elicited by endpoint cursor feedback (n=20). **A**, Grand averaged ERPs recorded from electrode FCz. ERPs are aligned to endpoint cursor feedback presentation (0 ms: vertical gray line). Horizontal grey bar indicates time window for FRN/RP analysis (267-317ms). Trials were selected for feedback rotation (0°, ±2°, or ±4°) and drug condition (levodopa or placebo) for ERP averaging. **B**, ERP amplitude during the FRN/RP time window. Individual participants’ data show amplitude following unrotated feedback as well as feedback rotated by ±2°, and ±4°. Differences in ERP amplitude between rotated and unrotated feedback are also shown for each participant. Boxplots indicate the median (circular markers), the interquartile range (thick bars) and the range (thin lines). **C**, Trial averaged ERPs recorded from electrode Pz. Horizontal grey bars indicate time window for P300 analysis (292-342 ms). **D**, ERP amplitudes during the P300 time windows, as in B.

P300: ERPs elicited by endpoint cursor feedback at electrode Pz are shown in Figure 6c. We analyzed the P300 by submitting the average ERP amplitude at electrode Pz between 292-342 ms to repeated measures ANOVAs (figure 6d). We did not find reliable main effects of drug (F(1,19) = 0.43, p = 0.52, 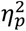=0.02), or feedback rotation (F(2,38) = 1.31, p = 0.28 (Greenhouse-Geisser corrected), 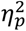= 0.06). We did observe a reliable drug*rotation interaction effect (F(2,38) = 7.46, p = 2.24e-3 (Greenhouse-Geisser corrected), 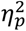= 0.28). Simple main effects revealed a reliable main effect of rotation in the placebo (F(2,38) = 5.72, p=6.72e-3) but not the levodopa (F(2,38) = 0.51, p=0.60) condition on P300 amplitude.

## Experiment 2

### Control measures

Participants’ judgments as to whether they received placebo or drug was near chance level (52.63%) and only 13.16% of participants responded that they thought they had received the drug. The values for heart rate, blood pressure, and alertness are reported in Table 2 for both the placebo and levodopa groups at the beginning and end of each experimental session. There were no reliable differences between the levodopa and placebo conditions in the percent change of heart rate (t(36) = -1.09, p=0.282, 95%CI for difference = [-0.10 0.03], BF = 0.5, posterior δ: median = - 0.273, 95%CI = [-0.875 0.284]), diastolic blood pressure (t(36) = 1.37, p=0.18, 95%CI for difference = [-0.02 0.11], BF = 0.65, posterior δ: median = 0.346, 95%CI = [-0.218 0.960]), systolic blood pressure (t(36) = 1.37, p=0.18, 95%CI for difference = [-.02 0.09], BF = 0.65, posterior δ: median = 0.346, 95%CI = [-0.218 0.960]), or alertness (t(36) = - 0.88, p=0.39, 95%CI for difference = [-0.95 0.38], BF = 0.43, posterior δ: median = - 0.218, 95%CI = [-0.810 0.337]). There was also no reliable difference between peak movement velocity between the levodopa and placebo groups (t(36) = -0.09, p=0.93, 95%CI for difference = [-0.01 9.94e-3], BF = 0.32, posterior δ: median = -0.021, 95%CI = [-0.585 0.539]).

**Table 3:**
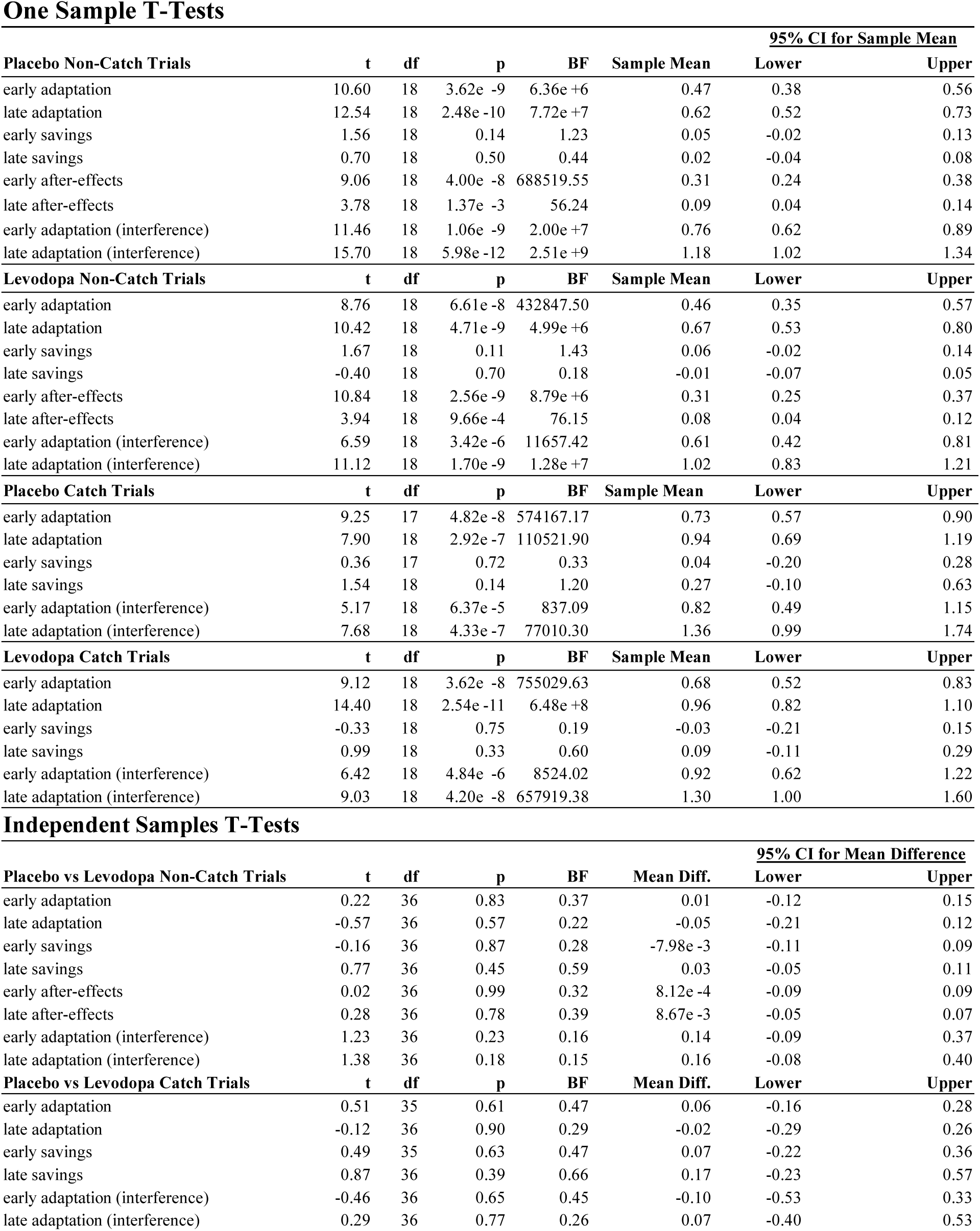
Statistical results for experiment 2. In one-sample T-Tests, the null hypothesis was that the mean was equal to zero. T, T-statistic. DF, degrees of freedom. P, P-value. BF, Bayes factor in favor of the alternative hypothesis. 95% CI, frequentist confidence interval. Mean differences are computed as placebo-levodopa. Bayes factors were computed using one-tailed default priors for the alternative hypothesis. In all one-sample T-Tests, the alternative hypothesis was that the population mean is greater than zero. For independent T-Tests, the alternative hypothesis stated that adaptation with interference would be greater in the levodopa group than the placebo group. For all other independent T-tests, the alternative hypothesis stated that the measure of interest would be smaller in the levodopa group than the placebo group.

#### Force field adaptation results

In each trial, we measured the perpendicular deviation (PD) of the reach trajectory at peak tangential velocity. PD data from throughout each force field and null field block, excluding catch trials, are shown in Figure 7. PD data from catch trials are shown in Figure 8. We computed contrasts to test for adaptation, savings, after-effects, and learning with interference in both the early (bins 1-5) and late (bins 6-12) periods following perturbation onset (Figure 9). We tested whether these effects are different from zero using 1-sample T-Tests for both the levodopa and placebo groups. We tested for differences between the levodopa and placebo groups using independent sample T-Tests. Detailed statistical results are shown in Table 3.

**Figure 7.**
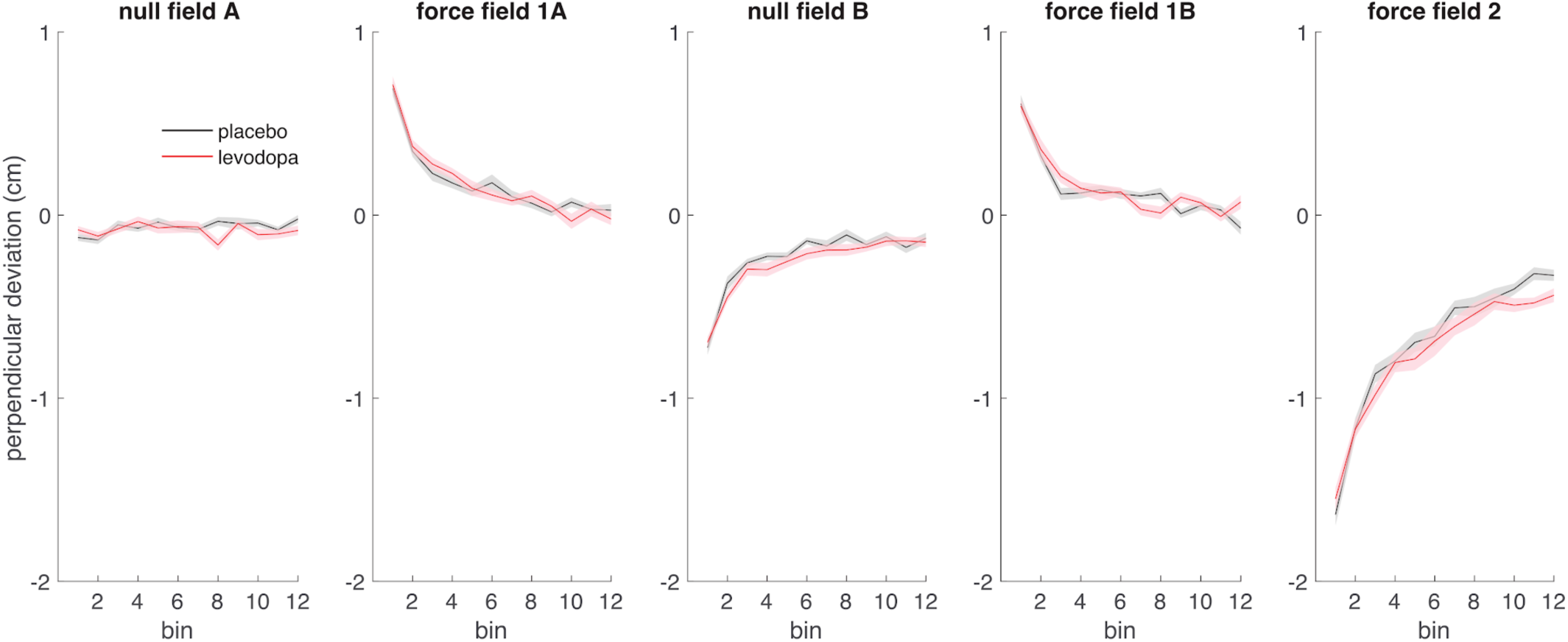
Perpendicular deviation of reach trajectory during non-catch trial reaches. Average perpendicular deviation of the hand trajectory within bins consisting of 8 trials each is shown in cm (Shaded region: ± SEM). The placebo condition is shown in black (n=19), and the levodopa condition is shown in red (n=19). Perpendicular deviation was measured on each trial at peak tangential velocity. Trials 6, 24, 35, 50, 71, and 91 of each block were catch trials, and were excluded from the corresponding bins. In *null field A* and *null field B,* the robot did not apply external forces to the hand during reaches. In *force field 1A* and *force field 1B,* participants made reaches in a clockwise force field. In *force field 2* participants made reaches in a counterclockwise force field.

**Figure 8.**
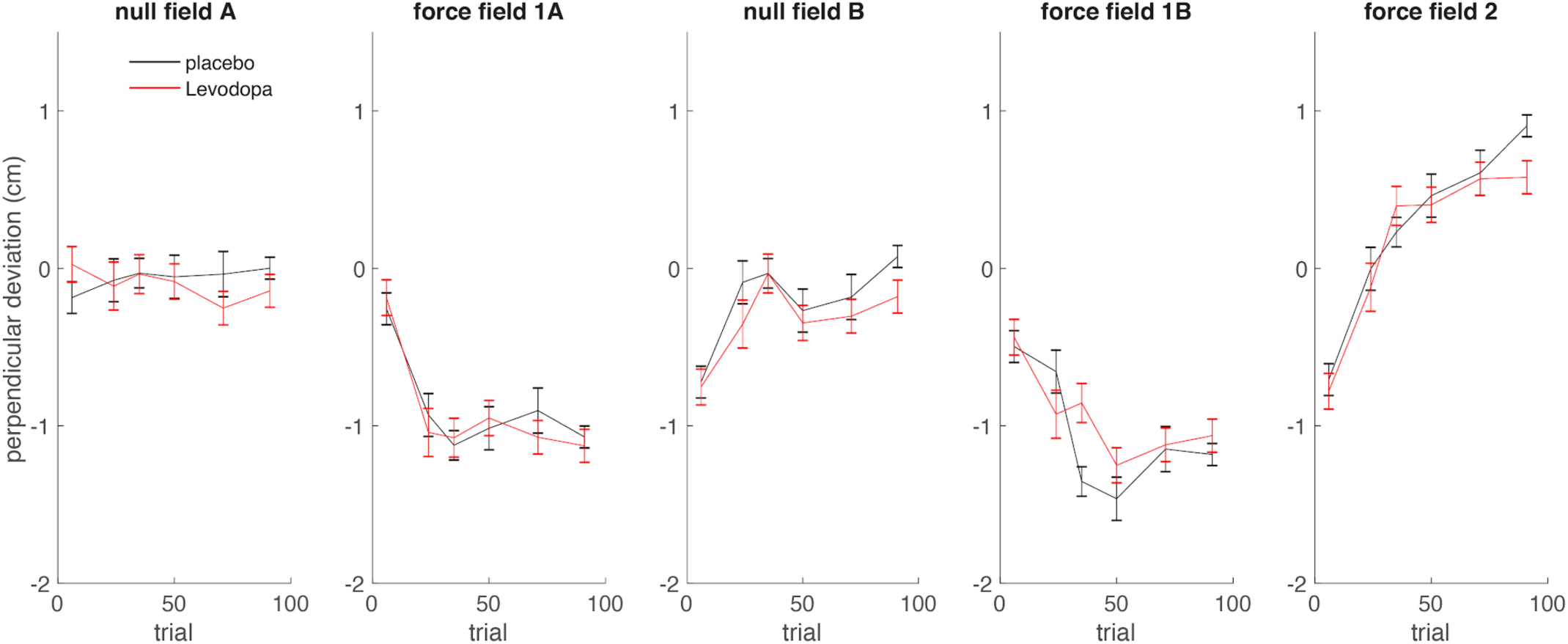
Perpendicular deviation of reach trajectory during catch trials. Perpendicular deviation of the hand trajectory, measured at peak tangential velocity, is shown in cm (Error bars: ± SEM). The placebo condition is shown in black (n=19), and the levodopa condition is shown in red (n=19). Catch trials occurred on trials 6, 24, 35, 50, 71, and 91 of each block. In *null field A* and *null field B,* the robot did not apply external forces to the hand during reaches. In *force field 1A* and *force field 1B,* participants made reaches in a clockwise force field. In *force field 2* participants made reaches in a counterclockwise force field.

**Figure 9.**
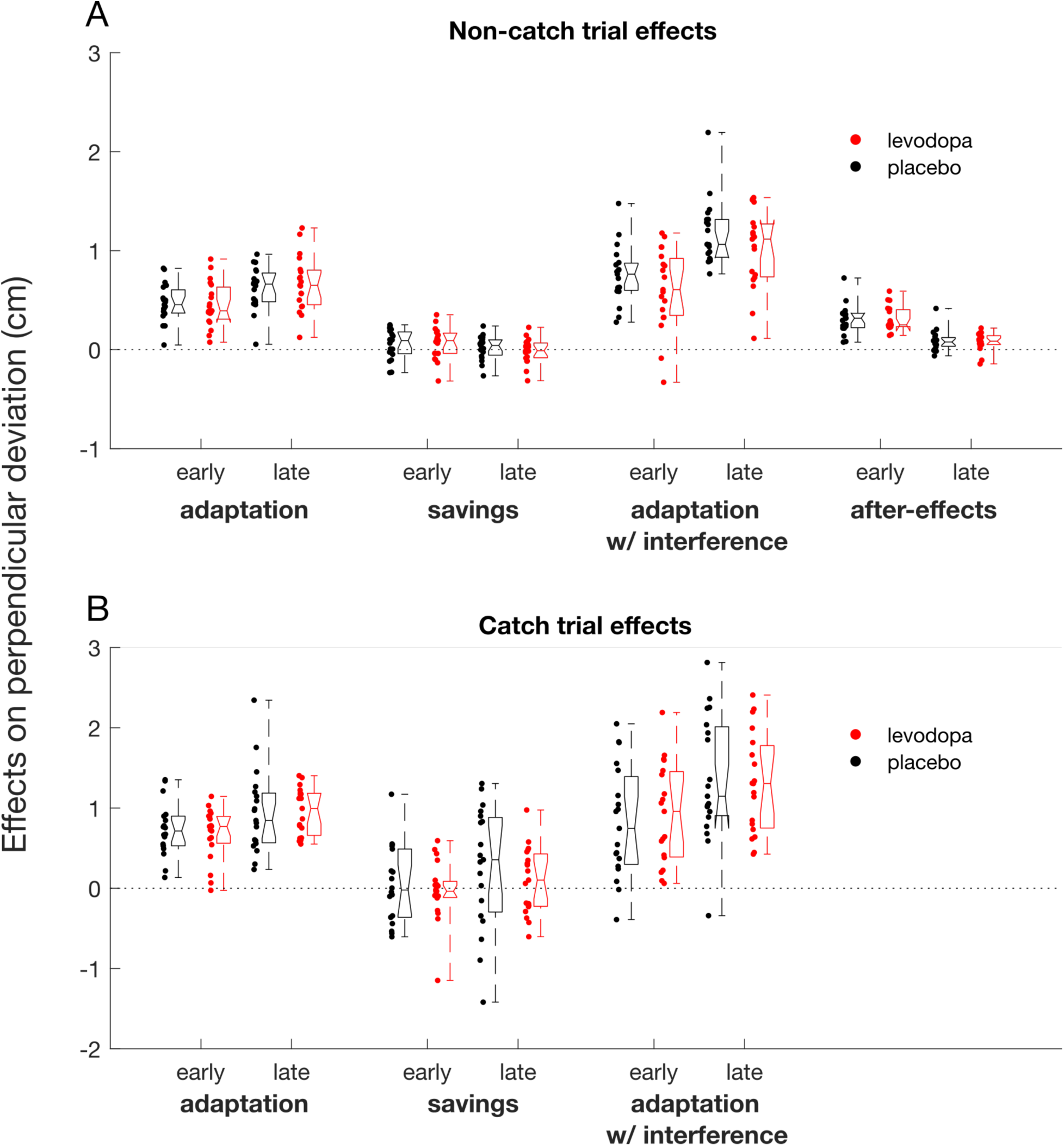
Adaptation effects in non-catch trials **A**, and catch trials **B**. Data points show effects for individual participants, box plots show the median, interquartile range, and full range. Effects are contrasts computed using perpendicular deviation (PD) of reach trajectory (cm), such that zero corresponds to no effect. Adaptation: change in PD during FF1a. Savings: difference in PD between FF1a and FF1b. After-effects: difference in PD between NFb and baseline from NFa. Adaptation w/ interference: change in PD during FF2.

#### Adaptation

##### Non-catch trials

Early adaptation was greater than zero in both the placebo (p=3.62e-9, BF=6.36e+6) and levodopa conditions (p=6.61e-8, BF=432848). We also observed reliable late adaptation for both the placebo (p=2.48e-10, BF=7.72e+7) and levodopa (p=4.71e-9, BF=4.99e+6) conditions. We did not observe a reliable difference between drug conditions for either early (p=0.83, BF=0.37) or late (p=0.57, BF=0.22) adaptation.

##### Catch trials

Early adaptation was greater than zero in both the placebo (p=4.82e-8, BF=574167) and levodopa (p=3.62e-8, BF=755029) conditions. We observed reliable late adaptation in both the placebo (p=2.92e-7, BF=110522) and levodopa (p=2.54e-11, BF=6.48e +8) conditions. There was no reliable difference between drug conditions for either early (p=0.61, BF=0.47), or late (p=0.90, BF=0.29) adaptation.

#### Savings

##### Non-catch trials

Our analyses yielded inconclusive evidence in favor of the hypothesized effect of savings for early adaptation for both the placebo (p=0.14, BF=1.23) and levodopa (p=0.11, BF=1.43) conditions. In the late period of adaptation, Non-catch trials provided inconclusive evidence against the hypothesized effect of savings following placebo (p=0.50, BF=0.44), and moderate evidence against the hypothesized effect of savings following levodopa (p=0.70, BF=0.18). There was moderate evidence against the hypothesis that savings would be reduced by levodopa in early adaptation (p=0.87, BF=0.28), and inconclusive evidence that savings would be reduced in late adaptation (p=0.45, BF=0.59).

##### Catch trials

There was moderate evidence against the hypothesized effects of savings for early adaptation following both placebo (p=0.72, BF=0.33) and levodopa (p=0.75, BF=0.19). Evidence for savings in late adaptation was inconclusive following both placebo (p=0.14, BF=1.20) and levodopa (p=0.33, BF=0.60). There was inconclusive evidence against the hypothesis that levodopa would reduce savings for both early (p=0.63, BF=0.47) and late (p=0.39, BF=0.66) adaptation.

#### After-Effects

##### Non-catch trials

We observed reliable after-effects in the early portion of NFb following adaptation in both the placebo (p=4.00e-8, BF=688519.55) and levodopa (p=2.56e-9, BF=8.79e+6) conditions. We also observed reliable after-effects extending to the later period of NFb after both placebo (p=1.37e-3, BF=56.24) and levodopa (p=9.66e -4, BF=76.15). We observed no reliable evidence that levodopa impaired after-effects in either the early (p=0.99, BF=0.32) or late (p=0.78, BF=0.39) periods.

#### Adaptation with interference

##### Non-catch trials

Early adaptation following exposure to an opposing force field was reliably greater than zero in both the placebo (p=1.06e-9, BF=2.00e+7) and levodopa (p=3.42e -6, BF=11657.42) conditions. We also observed reliable late adaptation in both the placebo (p=5.98e -12, BF=2.51e +9) and levodopa (p=1.70e -9, BF=1.28e +7) conditions. We observed moderate evidence against the hypothesized effect that levodopa would result in improved adaptation with interference in both the early (p=0.23, BF=0.16) and late (p=0.18, BF=0.15) periods.

##### Catch trials

Early adaptation following exposure to an opposing force field was reliably greater than zero in both the placebo (p=6.37e-5, BF=837.09) and levodopa (p=4.84e-6, BF=8524.02) conditions. We also observed reliable late adaptation in both the placebo (p=4.33e-7, BF=77010.30) and levodopa (p=4.20e-8, BF=657919.38) conditions. We observed inconclusive evidence against the hypothesis that levodopa would result in improved adaptation with interference in the early period (p=0.65, BF = 0.45), and moderate evidence in the late period (p = 0.77, BF = 0.26).

## Discussion

We tested for effects of levodopa, a dopamine precursor, in three different motor adaptation tasks across two experiments. In the first experiment we recorded EEG during a reward-based motor adaptation task and a sensory error-based visuomotor rotation (VMR) adaptation task. In the second experiment, we used a force field adaptation paradigm to test for effects of levodopa on initial adaptation, savings, and anterograde interference. We hypothesized that levodopa would selectively impair neural and behavioral responses to reinforcement feedback in the reward-based learning task as well as savings and interference. However, the only reliable influence of levodopa was in modulating the effect of visuomotor rotation on the P300 event-related potential component.

### Visuomotor rotation task

During the VMR task included in experiment 1, a cursor appeared at the endpoint of each reach to represent the position of the hand, and this feedback was perturbed through random rotations. We observed robust trial-by-trial adaptation to these perturbations. We did not find evidence that adaptation was affected by levodopa. This was expected, as trial-by-trial error correction induced by relatively small visuomotor rotations is thought to be driven primarily by sensory error-based learning mechanisms as opposed to dopaminergic reinforcement learning circuits (Diedrichsen et al., 2005; Ito, 2000; Krakauer et al., 2004; Tanaka et al., 2009; Jordan A. Taylor et al., 2010; Wong et al., 2019).

It has previously been shown that visuomotor rotation increases the amplitude of the P300 ERP component, a centro-parietal ERP deflection peaking approximately 300- 400ms following feedback presentation (Aziz et al., 2020; MacLean et al., 2015; Palidis et al., 2019). In the present study, we observed an interaction effect between feedback rotation and drug condition on the P300 amplitude. P300 amplitude increased in response to visuomotor rotations in the placebo condition but not in the levodopa condition. This result replicates previous findings that visuomotor rotations increase the amplitude of P300 responses to feedback, and additionally suggests that this effect is dependent on dopaminergic signaling. The modulation of P300 amplitude by sensory error is clearly not essential for adaptation, as disruption of this effect by levodopa did not correspond with any behavioral changes. Previous findings have also suggested a relationship between dopamine function and the P300 response, however the neural mechanisms and functional significance of the P300 in relation to motor adaptation remain unclear (Chu et al., 2018; Hansenne et al., 1995; Mulert et al., 2006; Noble et al., 1994; Sohn et al., 1998; Stanzione et al., 1990, 1991; Takeshita & Ogura, 1994). Variants of the P300 are elicited by many types of task-relevant stimuli, and have been localized to diffuse cortical areas including parietal, frontal, and motor regions, which have been implicated in processing prediction error (Bledowski et al., 2004; Calhoun et al., 2006; Johnson et al., 2019; Li et al., 2009; Mantini et al., 2009; Polich, 2007; Ragazzoni et al., 2019; Sabeti et al., 2016; Soltani & Knight, 2000). We observed a similar interaction effect between rotation and drug condition in recordings from electrode FCz during the FRN/RP time window. This appeared to be largely attributable to the P300 effect described above, as the time windows were mostly overlapping and the P300 was clearly measured at FCz as well.

### Reward learning task

Participants adapted reliably to manipulations of binary reinforcement feedback intended to produce either progressively clockwise or counterclockwise reach angles. However, we found no effects of levodopa on adaptation. One explanation of our findings is that the behavioral and neural processes measured in the current study do not depend on dopaminergic reward learning mechanisms. Another possibility is that the drug manipulation was not sufficiently powerful to disrupt these processes. The former interpretation depends on previous findings that levodopa impairs cognitive forms of reward learning using the same drug administration protocols in similar populations. However, the current study is limited by the lack of a positive control task demonstrating known behavioral effects of levodopa. Quattrocchi et al. (2018) found no effect of levodopa or a dopamine antagonist haloperidol on modulation of sensory error-based learning by additional reinforcement feedback. Holland et al. (2019) found no association between dopamine-related gene polymorphisms on adaptation through binary reinforcement feedback in a task similar to that used in the current study. Together, these findings suggest that reward-based motor adaptation may not rely on dopamine function, or at least that additional mechanisms may compensate for differences in dopamine function.

The “dopamine overdose” hypothesis states that dopaminergic medications such as levodopa might disrupt learning processes mediated by the ventral striatum by overstimulating dopamine signaling in this brain region. The ventral striatum may specifically mediate stimulus-based reinforcement learning, while action-based reinforcement learning in the current study may be subserved by the dorsal striatum (Rothenhoefer et al., 2017). Furthermore, levodopa may specifically impair learning from unfavorable outcomes as opposed to rewarding outcomes (Cools et al., 2006, 2007; Frank et al., 2004; Vo et al., 2018). Non-reward outcomes in the current task may not contribute significantly to learning as they do not instruct the correct response, unlike in binary response tasks.

Another important distinction is between model-free and model-based reinforcement learning processes (Babayan et al., 2018; Daw et al., 2011; Deserno et al., 2015; Dolan & Dayan, 2013; Doll et al., 2016; Gardner et al., 2018; Gläscher et al., 2010; Russek et al., 2017; Sambrook et al., 2018; Shahar et al., 2019; Sharp et al., 2016). Model-free reinforcement learning is characterized by reinforcement of simple stimulus-response associations that facilitate habitual, reflexive responding. Model-based learning allows for flexible planning according to a mental representation of the task, and can be limited by working memory processes. Levodopa has been shown to impair reward-based learning in healthy controls and people with Parkinson’s disease, but to improve model-based learning and related cognitive functions such as working memory, cognitive flexibility, and attention (Beato et al., 2008; R. Cools et al., 2001; Roshan Cools et al., 2003; Cooper et al., 1992; Costa et al., 2003; Kulisevsky, 2000; Lange et al., 1992; Lewis et al., 2005; Marini et al., 2003; Moustafa et al., 2008; Sharp et al., 2016; Torta et al., 2009; Wunderlich et al., 2012). It is possible that “dopamine overdose” by levodopa selectively impairs model-free learning. It may be that reward-based motor adaptation in the current study relies on processes other than model-free learning that are not affected by levodopa. Reward-based motor adaptation tasks similar to that in the current study have been shown to primarily involve strategic aiming that can be influenced by explicit instructions and cognitive load, characteristics that are inconsistent with model-free learning (Codol et al., 2018; Holland et al., 2018).

We also analyzed the variability of trial-by-trial changes in reach angle as a function of reward outcomes. Reward related modulation of motor variability has been shown to be impaired in medicated Parkinson’s disease in a very similar task (Pekny et al., 2015). We hypothesized that this effect may be due to side-effects of dopaminergic medication, and that we would observe similar impairments in healthy participants after levodopa administration. However, we observed no effect of levodopa on reward-related modulation of motor variability. Reward-based modulation of exploratory variance may therefore not depend on the ventral striatum, which is relatively spared in early stage Parkinson’s disease and therefore vulnerable to “dopamine overdose” in patients and healthy controls alike. Instead, it may depend on the dorsal striatum, which is more closely related to movement planning and is primarily impacted by early stage Parkinson’s disease.

Reinforcement feedback elicited a very reliable FRN/RP ERP component. Meta analyses have shown that the FRN/RP encodes a quantitative reward prediction error across multiple different tasks (Sambrook & Goslin, 2015; Walsh & Anderson, 2012). Reports have linked the FRN/RP signal to behavioral adjustments in response to feedback (Arbel et al., 2013; Frank et al., 2005; Holroyd & Krigolson, 2007; van der Helden et al., 2010). These findings support a prominent theory purporting that the FRN/RP is a reflection of reinforcement learning processes in the anterior cingulate cortex driven by phasic dopamine reward prediction error signals (Holroyd & Coles, 2002; Walsh & Anderson, 2012). Contrary to our hypothesis, we observed no effects of levodopa on the FRN/RP in response to reinforcement feedback. Previous studies have supported a link between dopamine and the FRN/RP, although results have been mixed. FRN/RP amplitude has been shown to be impaired in Parkinson’s disease patients with apathy (Martínez-Horta et al., 2014). Brown et al. (2020) found that the reward positivity was impaired in Parkinson’s disease patients relative to controls ON levodopa but not OFF levodopa, consistent with the dopamine overdose hypothesis. In healthy participants, the dopamine antagonist haloperidol has shown mixed results in reducing the amplitude of the reward positivity (Forster et al., 2017; Schutte et al., 2020). Mueller et al. (2014) found that the D2 receptor dopamine antagonist sulpiride had opposite effects on FRN/RP amplitude depending on a genotype variant that regulates prefrontal dopamine levels. They suggested a u-shaped relationship between dopamine release in the prefrontal cortex and FRN/RP amplitude mediated by the balance between D1 and D2 receptor activation. Because the effect of dopamine manipulation on the FRN/RP seems to depend on genetic differences in baseline dopamine release, one possibility is that levodopa in the current study had inconsistent effects on different subgroups of participants that cancelled each other in the group average.

### Force field adaptation task

Participants reliably adapted to the clockwise force field imposed in blocks FF1a and FF1b, and we found no evidence that adaptation was affected by levodopa. This was expected as force field adaptation is thought to rely primarily on sensory error-based learning mechanisms involving the cerebellum. Savings and interference effects have been accounted for by an additional process involving operant reinforcement of adapted motor commands upon repetition of successful reaches (Huang et al., 2011). These distinctions are supported by findings that cerebellar degeneration impairs force field adaptation while Parkinson’s disease patients are spared in initial adaptation but display deficient savings and interference (Bédard & Sanes, 2011; Leow et al., 2012, 2013; Maschke et al., 2004; Jordan A. Taylor et al., 2010). Thus, we hypothesized that dopaminergic perturbation by levodopa would impair savings and interference while leaving initial adaptation intact. We found no effect of levodopa on savings or interference. Impaired savings may therefore be a specific effect of Parkinson’s disease as opposed to a side-effect of levodopa. This is consistent with the findings of Marinelli et al. (2009), who observed a lack of savings effects in drug-naive and off-medication PD patients. An important limitation is that our experimental protocol was likely insufficient to produce savings or interference even in the control group, as we observed unreliable evidence of savings overall. Savings and interference have been shown to depend on sufficient repetition of the adapted movements to produce reinforcement of the adapted movements (Huang et al., 2011; Orban de Xivry & Lefèvre, 2015). Because the current study involved reaches to 8 different targets, repetition and each individual target was limited relative to single target experiments.

### Conclusions

As we expected, sensory error-based motor adaptation induced by visuomotor rotations and force field perturbations was not vulnerable to disruption of dopamine signaling by levodopa. This supports the notion that sensory error-based learning is driven by circuits involving cerebellar and sensorimotor cortex distinct from dopaminergic reinforcement learning mechanisms. Contrary to our hypotheses, we did not observe effects of levodopa on reward-based motor learning or the FRN/RP ERP component, which have both been theorized to depend on dopaminergic signaling of reward prediction error. The dopamine overdose hypothesis suggests that levodopa impairs stimulus-response reinforcement learning processes in the ventral striatum. Reward-based motor adaptation may instead depend on distinct reinforcement learning circuits that are not disrupted by levodopa such as cortical reward learning mechanisms or dopaminergic projections to the dorsal striatum.

